# ABI5-mediated ABA signaling enhances aliphatic glucosinolates biosynthesis by transcriptionally suppressing BR signaling factors in *Arabidopsis*

**DOI:** 10.64898/2026.03.10.710799

**Authors:** Dasom Choi, Hyunjoon Kim, Dong-Hwan Kim

**Affiliations:** Department of Plant Science and Technology, Chung-Ang University, Anseong, 17546, Republic of Korea

**Author notes:** **Correspondence to:** Dong-Hwan Kim, Department of Plant Science and Technology, Chung-Ang University, Anseong, Gyeonggi-do, 17546, Republic of Korea.

**Keywords:** Glucosinolates, Brassinosteroids, Abscisic acid, ABI5, BZR1, Histone modification

## Abstract

Brassinosteroids (BRs) are key regulators of plant growth and have been implicated in suppressing glucosinolates (GSLs) biosynthesis in Brassicaceae species, including *Arabidopsis thaliana*. However, the molecular mechanism linking BR signaling to transcriptional control of the aliphatic GSL pathway remains unclear. Here, we provide genetic and molecular evidence that the BR-responsive transcription factor BZR1 negatively regulates aliphatic GSL biosynthesis through interaction with TOPLESS (TPL) family corepressors and the histone deacetylase HISTONE DEACETYLASE 19 (HDA19). The gain-of-function mutant *bzr1-1D* exhibited reduced accumulation of aliphatic glucosinolates (GSLs), whereas disruption of *TPL* family genes (*tpl, tpr1, tpr4*) or *HDA19* resulted in elevated GSL levels, supporting a repressive role for the BZR1-associated complex in the regulation of GSL biosynthetic gene expression. Furthermore, we uncover a functional connection between ABA and BR signaling in this process. The ABA-responsive transcription factor ABI5 reduces the expression of *UBP12* and *UBP13*, which encode deubiquitinases known to influence BZR1 stability. Genetic and transcriptional analyses indicate that ABI5-mediated attenuation of the UBP12/13-BZR1 pathway contributes to enhanced aliphatic GSL accumulation under ABA treatment. Collectively, our findings delineate a regulatory framework linking ABA and BR signaling pathways and suggest that modulation of the UBP12/13-BZR1 module by ABI5 integrates growth and defense responses by fine-tuning aliphatic GSLs biosynthesis in *Arabidopsis*.

**One sentence summary:** an ABA signaling factor, ABI5 acts to suppress expression of brassinosteroids (BR) hormone signaling genes like UBP12, UBP13, and BZR1 to promote biosynthesis of defensive secondary metabolites, glucosinolates (GSLs) in *Arabidopsis*.

## INTRODUCTION

Plant orchestrates a complex defense mechanism upon environmental cues against biotic and abiotic stress. Belonging to secondary metabolites groups, glucosinolates (hereafter called as GSLs) play an important role in plant defense system of Brassicaceae family plants including *Arabidopsis thaliana*. GSLs is a group of nitrogen and sulfur-containing secondary metabolites whose degraded products are responsible for intrinsic flavors and pungent aroma (Heber, 2004, Grubb and Abel, 2006, Higdon et al., 2007). In addition, GSLs have healthy dietary effects to human and also play a role as cancer preventive compounds (Higdon et al., 2007, Heber, 2004). GSLs are mainly produced by *Brassicaceae* family plants including dicot model plant *Arabidopsis* but also present in other plants (Fahey et al., 2001). The structure and composition of GSLs vary in different tissues and ages, environmental conditions, and variety. More than 130 different types of GSLs have been identified in a variety of plants to date (Nguyen et al., 2020). Depending on the amino acid precursors, GSLs are categorized into three different major groups: aliphatic, indolic, and aromatic GSLs (Fahey et al., 2001, Halkier and Gershenzon, 2006). Biosynthetic process for GSLs production takes place in three major steps: (i) chain elongation; (ii) core structure formation, and (iii) secondary modification (Zang et al., 2009, Mitsui et al., 2015) (Fig. 1A). However, indolic GSLs does not undergo the chain elongation step (Petersen et al., 2019).

**Figure 1.**
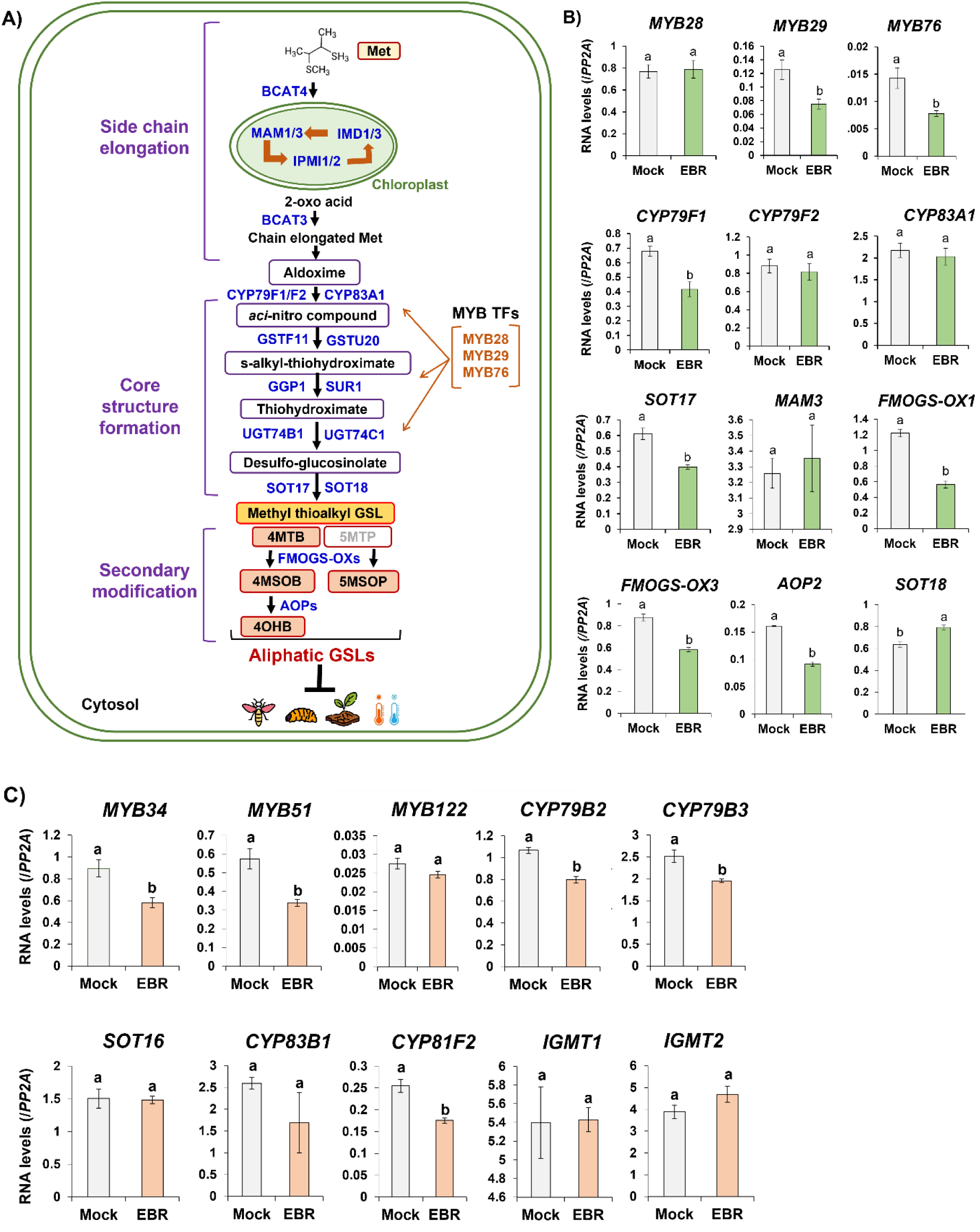
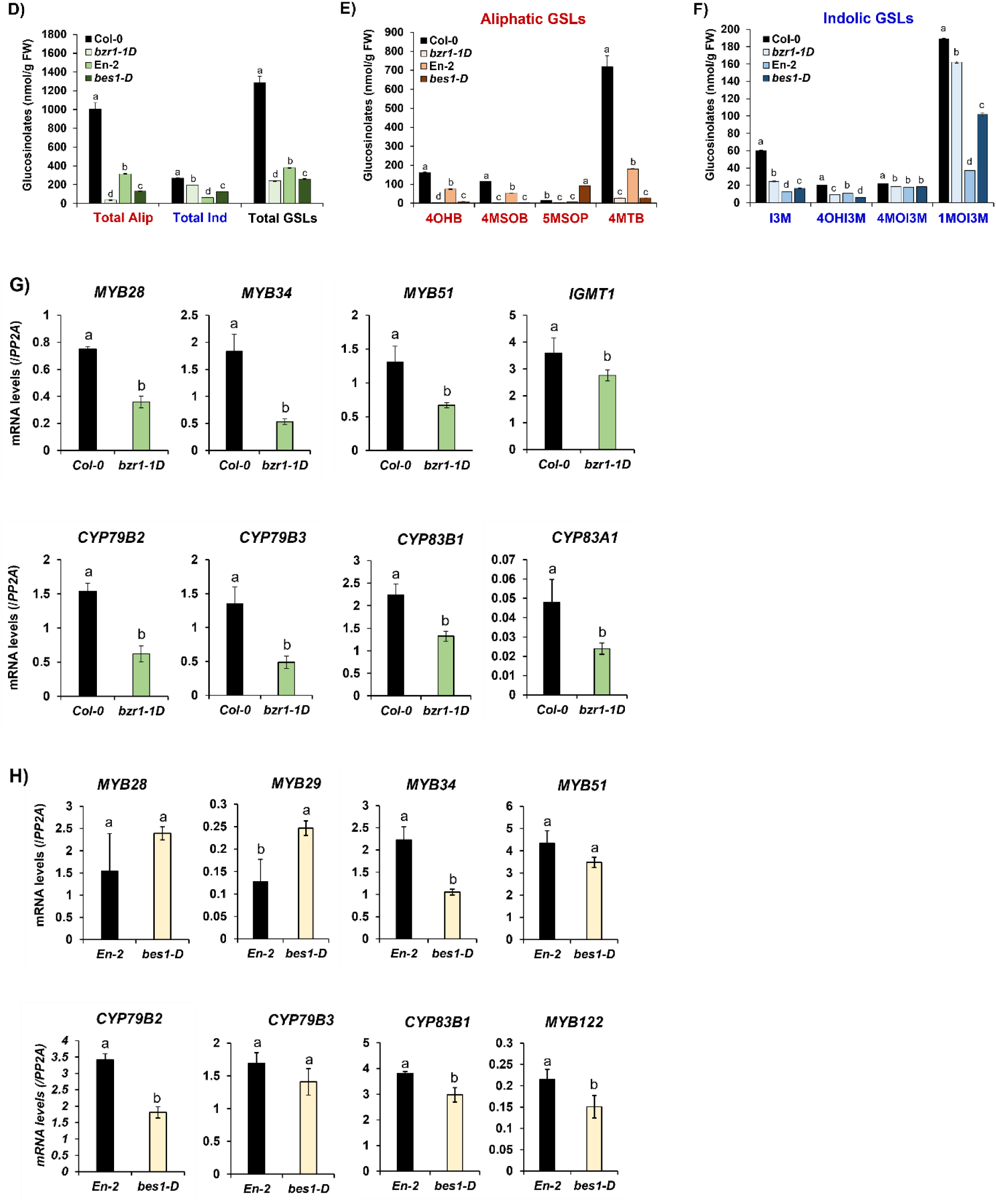
A) The schematic diagram showing aliphatic GSLs biosynthesis pathway in *Arabidopsis thaliana* model plant. Production of aliphatic GSL compounds requires three biosynthetic phases like 1) side-chain elongation, 2) Core structure formation, and 3) secondary modification (indicated with purple color letters). A handful of MYB transcription factors like MYB28, MYB29, and MYB76 (indicated with orange letters) were shown to control expression of biosynthetic pathway genes related to aliphatic GSL production. Produced aliphatic GSL compounds exerts a diversity of defensive roles against biotic (e.g. insect and larva) and abiotic (e.g. drought, cold etc.) challenges. The blue letters indicate the enzymes responsible for the conversion of the aliphatic compounds throughout the GSL biosynthetic process. **Abbreviations of catalytic enzymes**: METHYLTHIOALKYLMALATE SYNTHASE 1 (MAM1)/METHYLTHIOALKYMALATE SYNTHASE-LIKE (MAM3); ISOPROPYLMALATE DEHYDROGENASE 1 (IMD1)/IMD3; ISOPROPYLMALATE ISOMERASE 1 (IPMI1)/IPMI2; BRANCHED CHAIN AMINO ACID TRANSFERASE 3 (BCAT3); CYTOCHROME P450 79F1 (CYP79F1)/CYP79F2; GLUTATHIONE S-TRANSFERASE F11 (GSTF11); GLUTATHIONE S-TRANSFERASE TAU 20 (GSTU20); GAMMA-GLUTAMYL PEPTIDASE 1 (GGP1); SUPERROOT 1 (SUR1); UDP-GLUCOSYL TRANSFERASE 74B1 (UGT74B1); UDP-GLUCOSYL TRANSFERASE 74C1 (UGT74C1);SULFOTRANSFERASE 16 (SOT16); SULFOTRANSFERASE 17 (SOT17); SULFOTRANSFERASE 18 (SOT18); FLAVIN-MONOOXYGENASE GLUCOSINOLATE S-OXYGENASES (FMO GS-OXs); ALKENYL HYDROXALKYL PRODUCINGS (AOPs). **Abbreviations for aliphatic GSL compounds**: 4-hydroxybutyl (4OHB), 4-methylsulfinlybutyl (4MSOB), 5-methylsulfinylpentyl (5MSOP), and 4-methylthiobutyl (4MTB). **B)** Result of qRT-PCR analysis on 12 randomly selected aliphatic GSL pathway genes between the Mock (non-treated) and 24-Epibrassinolide (EBR)-treated samples. seven genes (*MYB29, MYB76, CYP79F1, SOT17, FMOGS-OX1, FMOGS-OX3,* and *AOP2*) out of 12 aliphatic GSL pathway genes were significantly downregulated by EBR and only *SOT18* was slightly upregulated in EBR-treated samples in comparison to the Mock control sample. **C)** Result of qRT-PCR analysis on 10 randomly selected indolic GSL pathway genes between the Mock (non-treated) and 24-Epibrassinolide (EBR)-treated samples. Five genes (*MYB34, MYB51, CYP79B2, CYP79B3*, and *CYP81F2*) out of 10 selected genes exhibited down-regulation by EBR treatment in comparison to the Mock control. **D)** Quantification of aliphatic and indolic GSL compounds between Col-0 and the *bzr1-1D* gain-of-function mutant or En-2 and the *bes1-D* (En-2) gain-of-function mutant seedlings. Amounts of total GSLs were calculated by combining amounts of aliphatic and indolic GSL compounds. Alip: aliphatic, Ind: indolic. **E)** Quantification of aliphatic GSL compounds between Col-0 and the *bzr1-1D* gain-of-function mutant or En-2 and the *bes1-D* (En-2) gain-of-function mutant seedlings. Amounts of total aliphatic GSLs were calculated by combining amounts of four individual aliphatic GSL compounds like 4OHB, 4MSOB, 5MSOP, and 4MTB. **F)** Quantification of indolic GSL compounds between Col-0 and the *bzr1-1D* gain-of-function mutant or En-2 and the *bes1-D* (En-2) gain-of-function mutant seedlings. Amounts of total indolic GSLs were calculated by combining amounts of four individual indolic GSL compounds like I3M, 4OHI3M, 4MOI3M, and 1MOI3M. **G)** Result of qRT-PCR analysis on eight aliphatic and indolic GSL pathway genes between Col-0 wild type and *bzr1-1D* mutant. **H)** Result of qRT-PCR analysis on eight aliphatic and indolic GSL pathway genes between En-2 wild type and *bes1-D* (En-2 background) mutant. **B)∼H)** Two-week-old seedlings grown under long day (16h light: 8h dark) at 22℃ were harvested at ZT4 and subjected for RT-qPCR analysis. One-way ANOVA with Tukey’s post-hoc test was applied to calculate the statistical significance, and significant difference was indicated in the figures by different letters (p < 0.05). Data were presented as means ± standard deviation (SD) of three biological replicates.

During last decades, genes for GSLs metabolism have been identified by intensive genetic and biochemical analyses, mainly using *Arabidopsis* plants. In case of aliphatic GSLs (Fig. 1A), it starts from the deamination of an amino acid (i.e. methionine) to a corresponding 2-oxo acid by the catalytic activity of the BRANCHED CHAIN AMINO ACID TRANSFERASE 4 (BCAT4) in the cytosol (Schuster et al., 2006). BILE ACID TRANSPORTER 5 (BAT5) transports the 2-oxo acid into the chloroplast for a sequential catalytic reactions. In Chloroplast, condensation, isomerization, and oxidative decarboxylation of 2-oxi acid occurs by METHYLTHIOALKYL MALATE SYNTHASE 1∼3 (MAM1∼3), ISOPROPYLMALATE ISOMERASES 1/2 (IPMI1/2), and ISOPROPYLMALATE DEHYDROGENASE 1/3 (IMD1/3), respectively (Petersen et al., 2019, Mitreiter and Gigolashvili, 2021). After a series of reactions in the chloroplast, the 2-oxo acid is returned back to the cytosol and undergoes the transamination by BRANCHED CHAIN AMINO ACID TRANSFERASE 3 (BCAT3), forming branched-chain amino acids (Halkier and Gershenzon, 2006).

Branched-chain amino acids from first step enters to the second step, “core structure formation” in the cytosol (Fig. 1A). Cytochrome P450 monooxygenase family proteins named as CYTOCHROME P450 79F1 MONOOXYGENASE (CYP79F1)/CYP79F2 converts the branched-chain amino acids into the aldoximes which are further oxidized to nitrile oxide compounds by the catalytic activity of CYTOCHROME P450 83A1 MONOOXYGENASE (CYP83A1). Next, this nitrile oxide compounds are conjugated with glutathione by the catalytic activity of GLUTATHIONE S-TRANSFERASE F11 (GSTF11) and GLUTATHIONE S-TRANSFERASE U20 (GSTU20), then cleaved via γ-GLUTAMYL PEPTIDASE 1 (GGP1) and C-S LYASE, SUPER ROOT 1 (SUR1) to generate thiohydroximates (Halkier and Gershenzon, 2006). In the last reaction, thiohydroximate is converted to desulfo-GSLs by the S-glycosylation by UDP-glucosyltransferase 74C1 (UGT74C1) (Sonderby et al., 2010). Desulfo-GSLs are further sulfated by two sulfotransferases (SOT17 and SOT18), finally forming the methylthioalkyl GSLs (Hirschmann et al., 2017). In the last step, “side-chain modification” of aliphatic GSLs, methylthioalkyl GSL compounds undergoes various side-chain modifications like oxidation, alkylation, methoxylation, and desaturation (Fig. 1A). For instance, methylthioalkyl GSLs are sulfur-oxygenated by flavin monooxygenases (FMO GS-OXs) to generate methylsulfinylalkyl GSLs. Methylsulfinylalkyl GSLs are further converted to alkenyl GSLs by the catalytic activity of 2-oxoglutarate-dependent dioxygenase like alkenylhydroxalkyl-producing 2 (AOP2) (Hansen et al., 2008).

Transcriptional regulators controlling GSLs metabolism have been also identified. Six MYB domain proteins in *Arabidopsis* play a crucial role in the expression of GSLs biosynthetic genes. A subgroup of MYB TFs like MYB28, MYB29, and MYB76 acts to positively regulate expression of aliphatic GSLs pathway genes such as *MAM1*, *MAM3*, *CYP79F1*, *CYP79F2*, *CYP83A1*, *SUR1*, *SOT17*, and *SOT18* (Hirai et al., 2007, Sonderby et al., 2010, Gigolashvili et al., 2007, Beekwilder et al., 2008) (Fig.1A).

GSLs metabolism is affected by a diversity of endogenous (i.e. phytohormones) and environmental cues (i.e. wounding, herbivore, pathogen attack) (Yan and Chen, 2007, Malhotra and Bisht, 2020). Among plant hormones, Jasmonates (JAs) are lipid-derived cyclopentanones including jasmonic acid (JA) and its various derivatives and plays essential roles in plant defense against a diversity of stresses by triggering downstream signaling cascade (Kazan, 2015). Most of all, three basic helix-loop-helix TFs, MYC2, MYC3 and MYC4 execute a major function in JA-mediated stress response in *Arabidopsis* plant. These MYCs were shown to bind to E-box (CANNTG)/G-box (CACGTG) sequences in the promoters of defense-related genes (Fernandez-Calvo et al., 2011). One microarray analysis using triple mutants of MYCs, *myc2;3;4* indicated that these homologous MYCs TFs are positively require for many of GSLs pathway genes, thus showing greatly reduced amounts of GSLs in mutants (Schweizer et al., 2013). In addition, MYCs physically associate with MYBs, thus forming MYBs-MYCs heterodimer to positively regulate expression of the GSL biosynthetic genes (Frerigmann et al., 2014, Schweizer et al., 2013). It indicates that JA has a positive impact on GSL biosynthesis.

In contrast to JA, another plant hormone, Brassinosteroids (BRs), also called as a growth hormone, play a crucial role in a diversity of plant growth and development (Manghwar et al., 2022, Planas-Riverola et al., 2019). BRs signaling cascade is triggered by the perception of active form of BRs by a membrane-anchored receptors like BRASSINOSTERIOID INSENSITIVE 1 (BRI1) and BRI1-ASSOCIATED KINASE 1(BAK1) (Clouse, 2011). Then, signal is transmitted to two plant-specific TFs, BRASSINAZOLE-RESISTANT1 (BZR1) and BRI1-EMS-SUPPRESSOR1 (BES1) to initiate downstream BRs-associated genes (Kono and Yin, 2020, Shi et al., 2022). Under BR-sufficient condition, BZR1 and BES1 are dephosphorylated, stabilized, and transcriptionally active, whereas phosphorylation by kinase, BRASSINOSTEROID-INSENSITIVE 2 (BIN2) promotes their inactivation through the ubiquitin–proteasome machinery (Zhu et al., 2017, Nolan et al., 2020). Recent studies revealed that the stability of BZR1/BES1 is also controlled by deubiquitination. In particular, the UBIQUITIN-SPECIFIC PROTEASE 12 (UBP12) and UBP13 directly interact with BES1 and BZR1 (Wang et al., 2013a, Park et al., 2022). This interaction leads to the deubiquitination of BES1 (and likely BZR1), thereby reinforcing BES1/BZR1-mediated BR signaling (Wang et al., 2013a, Park et al., 2022, Xiong et al., 2022). Loss of *UBP12/UBP13* leads to increased ubiquitination and reduced levels of BES1, resulting in impaired BR signaling. Conversely, overexpression of either UBP12 or UBP13 enhances BES1 stability and BR sensitivity. Collectively, these findings highlight UBP12 and UBP13 as key regulators that reinforce BR signaling by maintaining the stability of BZR1 and BES1.

In terms of regulation of GSL biosynthesis, BRs have a negative impact on production of GSLs (Wang et al., 2023, Guo et al., 2013). For example, treatment of 24-epibrassinolide (EBR) inhibited the biosynthesis of aliphatic and indolic GSLs. In addition, it was previously shown that BZR1 and BES1 play a negative role in the production of GSLs (Yin et al., 2002). The *bes1-D* and *bzr1-1D* mutants which carry dominant negative mutation in *BES1* and *BZR1*, respectively had substantially reduced amounts of GSLs in *Arabidopsis.* Furthermore, it was recently reported that BZR1 and BES1 physically interact with MYBs controlling GSL biosynthesis, thus exerting a negative impact on production of GSLs (Liao et al., 2020). Another recent study also demonstrated that dephosphorylated BZR1 directly binds to BRRE (BR responsive element) located in the *MYB51* and *MYB29* genes for the regulation of indolic and aliphatic GSL biosynthesis, respectively (Wang et al., 2023). These data indicate that BR signaling acts negatively on the biosynthesis of GSLs in Arabidopsis.

It was previously reported that a ABA signaling factor, ABA-INSENSITIVE 5 (ABI5) is positively involved in the regulation of GSL biosynthesis (Miao et al., 2016, Miao et al., 2013). It suggested that GSL biosynthesis is fine-tuned by delicate regulatory crosstalk between JA, ABA, and BR signaling. While the crosstalk between JA and BR signaling was previously evidenced (Frerigmann et al., 2014, Schweizer et al., 2013), it is still merely unveiled how ABA and BR antagonistically crosstalk to modulate the endogenous biosynthesis of GSLs. In this study, we found that an ABA signaling factor, ABI5 acts to negatively regulate expression of *UBP12* and *UBP13* which are responsible for deubiquitination of BZR1 and BES1, thus enhancing BR signaling. In addition, we also found that ABI5 directly modulate transcription of *BZR1* and *BES1*. Our study revealed that ABI5-mediated enhancement of GSL biosynthesis is achieved by the direct suppression of BR signaling module genes like *UBP12/13* and its downstream factors, *BZR1/BES1* in Arabidopsis.

## RESULTS

### BR signaling factors play a negative role in GSLs biosynthesis

Previous studies have suggested that brassinosteroids (BRs) negatively regulate glucosinolates (GSLs) biosynthesis in *Arabidopsis* (Guo et al., 2013, Wang et al., 2023). To further examine the effects of BR signaling on GSL biosynthesis under our experimental conditions, we treated *Arabidopsis thaliana* Col-0 seedlings with 24-epibrassinolide (EBR), a biologically active brassinosteroid, and compared the expression levels of GSL biosynthetic genes between control (non-treated) and EBR-treated samples. Because aliphatic and indolic GSLs represent the major classes of GSL compounds in *Arabidopsis* (Choi et al., 2024), we selected a representative set of 12 aliphatic and 10 indolic GSL biosynthetic genes and analyzed their transcript levels in response to EBR treatment. Among the aliphatic GSL pathway genes, seven out of twelve genes (58%) were significantly downregulated by EBR treatment, whereas only *SOT18* showed a slight upregulation compared to the control (Fig. 1B). Similarly, five out of the ten examined indolic GSL pathway genes exhibited reduced expression following EBR treatment (Fig. 1C), indicating an overall repressive effect of BR on indolic GSL gene expression. Taken together, these results demonstrate that BR signaling suppresses the transcription of both aliphatic and indolic GSL biosynthetic genes in *Arabidopsis*, consistent with previous reports (Guo et al., 2013, Wang et al., 2023).

### BR signaling factor, BES1 and BZR1 suppress GSL pathway genes

To investigate whether the BR signaling components, BZR1 and BES1 are involved in the regulation of glucosinolates (GSLs) biosynthesis, we quantified GSL contents in wild-type plants and gain-of-function mutants of these key BR signaling factors, *bzr1-1D* and *bes1-D*. Overall, the total aliphatic GSL content was markedly reduced in both gain-of-function mutants, *bzr1-1D* and *bes1-D*, compared with their respective wild-type controls, Col-0 and En-2 (Fig. 1D). In contrast, the total indolic GSL content was significantly decreased in *bzr1-1D* relative to Col-0, whereas *bes1-D* exhibited higher levels of total indolic GSLs than the En-2 wild type (Fig. 1D). These results suggest that BES1-mediated regulation of indolic GSL biosynthesis may be more dynamic and complex than BZR1-mediated regulation of aliphatic GSL biosynthesis. Nevertheless, the total GSL content, calculated as the sum of aliphatic and indolic GSLs, was substantially reduced in both *bzr1-1D* and *bes1-D* compared with their corresponding wild-type plants (Fig. 1D).

Using HPLC-based analysis, we identified a total of eight distinct glucosinolates (GSLs) compounds, comprising four aliphatic and four indolic GSLs (Fig. 1E-1F; Supplementary Fig. S1; Supplementary Table S1). All four aliphatic GSLs (4OHB, 4MSOB, 5MSOP, and 4MTB) were significantly reduced in the *bzr1-1D* mutant compared with Col-0 plants (Fig. 1E). In contrast, the *bes1-D* mutant displayed significantly decreased levels of three aliphatic GSLs (4OHB, 4MSOB, and 4MTB), whereas the level of 5MSOP was increased, suggesting that BES1 plays a regulatory role in aliphatic GSL biosynthesis that is largely similar to, but partially distinct from, that of BZR1.

In case of indolic GSLs, all four detected compounds (I3M, 4OHI3M, 4MOI3M, and 1MOI3M) were consistently lower in *bzr1-1D* than in Col-0 (Fig. 1F). In contrast, only 4OHI3M was significantly reduced in *bes1-D*, whereas the remaining indolic GSLs (I3M, 4MOI3M, and 1MOI3M) were unchanged or even elevated compared with En-2 (Fig. 1F). Together, these results indicate that BZR1 and BES1-mediated BR signaling exerts an overall negative effect on GSL biosynthesis in *Arabidopsis*, while influencing aliphatic and indolic GSLs in distinct manners.

Next, we examined whether BZR1 and BES1 modulate endogenous GSL levels through transcriptional regulation of GSL biosynthetic genes. Transcript levels of eight representative GSL pathway genes were analyzed in Col-0 and *bzr1-1D*, as well as in En-2 and *bes1-D*. All eight GSL pathway genes were significantly downregulated in *bzr1-1D* compared with Col-0 (Fig. 1G), consistent with the reduced accumulation of aliphatic GSLs observed in *bzr1-1D* (Fig. 1D and 1E). These results indicate that BZR1 functions as a strong transcriptional repressor of GSL biosynthetic genes. In contrast, *bes1-D* displayed more variable transcriptional responses among GSL pathway regulators (Fig. 1H). For example, *MYB34*, a key MYB transcription factor regulating indolic GSL biosynthesis, was downregulated in *bes1-D* relative to En-2. Expression of two other MYB transcription factor genes was not significantly altered, whereas *MYB29* was upregulated in *bes1-D* compared with En-2 (Fig. 1H). These transcriptional patterns are consistent with the differential effects of BES1 on individual aliphatic and indolic GSL compounds (Fig. 1E and 1F). Taken together, our results demonstrate that BZR1 broadly suppresses both aliphatic and indolic GSL biosynthesis through transcriptional repression of GSL pathway genes, whereas BES1 exerts a more complex and differential regulatory effect on aliphatic and indolic GSL biosynthesis in *Arabidopsis*.

### Association of BZR1 with a co-repressor module, TPL-HDA19 is required for the suppression of GSL pathway genes

Given the clear evidence that BZR1 negatively regulates both aliphatic and indolic GSLs biosynthesis, we subsequently focused on BZR1 to further elucidate the molecular mechanisms underlying BZR1-mediated regulation of GSL biosynthesis in *Arabidopsis*. Previous studies have shown that BZR1 interacts with the histone deacetylase HDA19 and the co-repressor TOPLESS (TPL) to regulate the expression of downstream BR-responsive genes (Oh et al., 2014). This led us to examine whether BZR1 functions in conjunction with the TPL–HDA19 co-repressor complex to negatively regulate GSL biosynthesis. To test this possibility, we quantified the levels of aliphatic and indolic GSLs in Col-0, the *tpl* knockout mutant, and the *tpl;tpr1;tpr4* triple mutant, which is defective in TPL and its paralogs, TPR1 and TPR4. The total aliphatic GSL content was significantly increased in both *tpl* and *tpl;tpr1;tpr4* mutants compared with Col-0 (Fig. 2A). Notably, the extent of aliphatic GSL accumulation was comparable between the two mutants, suggesting that TPR1 and TPR4 may not play major roles in the regulation of aliphatic GSL biosynthesis in *Arabidopsis*.

**Figure 2.**
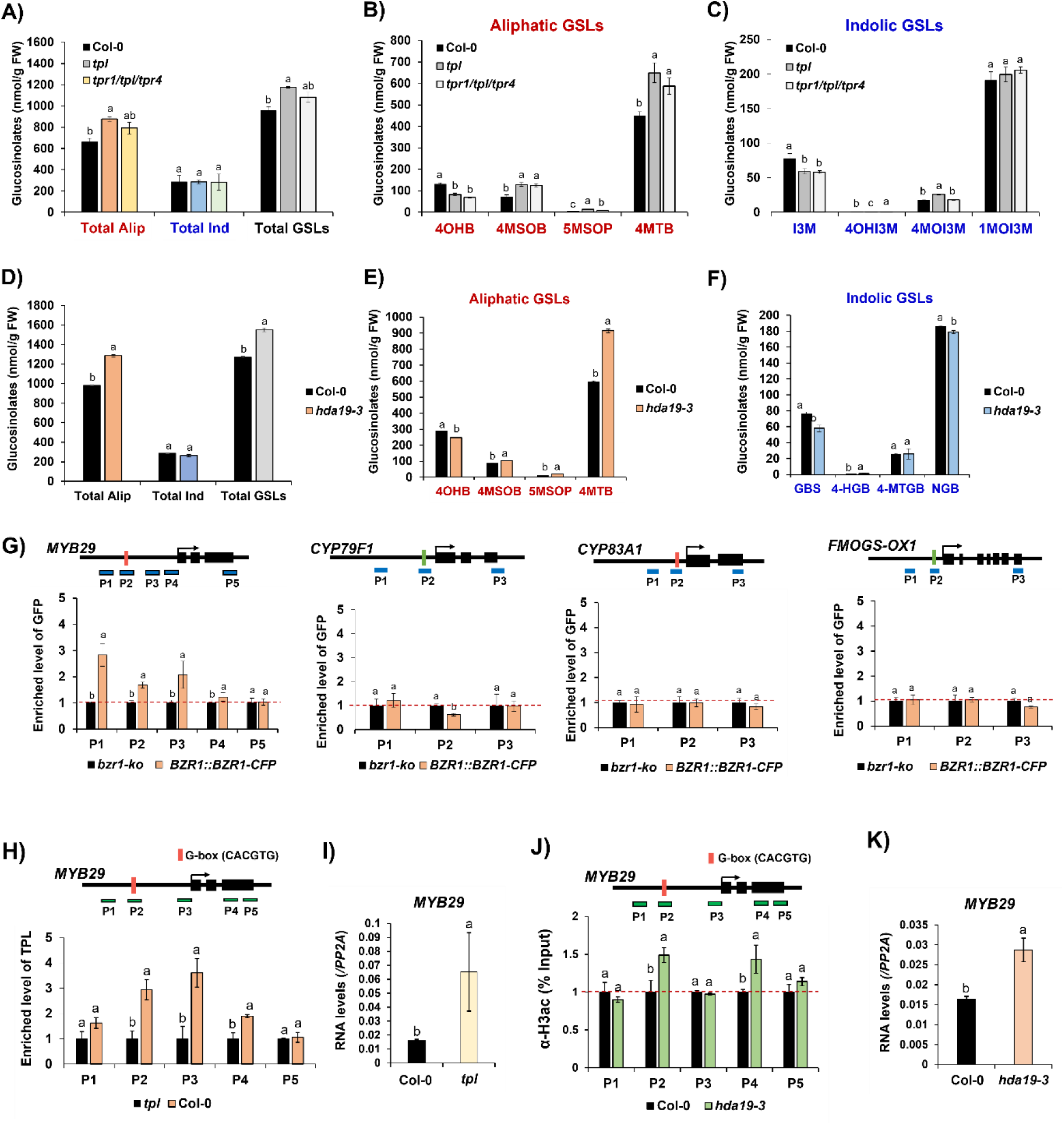
TPL(R)-HDA19 co-repressor complex is involved in the suppression of aliphatic GSLs production via histone deacetylation of *MYB29*. **A)** Quantification of aliphatic and indolic GSL compounds between Col-0 and the *tpl,* a knock-out mutant of *TPL,* and the *tpl/tpr1/tpr4,* a triple mutant of *TPL, TPR1, and TPR4*. Amounts of total GSLs were calculated by combining amounts of aliphatic and indolic GSL compounds. Amount of total aliphatic GSLs were substantially higher in both *tpl* and *tpr1/tpl/tpr4* mutants compared to that of Col-0. Meanwile, amount of total indolic GSLs did not show significant differences between Col-0, *tpl*, and *tpr1/tpl/tpr4* mutants. Alip: aliphatic, Ind: indolic. **B)** Quantification of four different aliphatic GSL compounds between Col-0, *tpl*, and *tpr1/tpl/tpr4* mutant seedlings. While three aliphatic GSL compounds like 4MSOB, 5MSOP, and 4MTB were significantly higher in both *tpl* and *tpl/tpr1/tpr4* mutants compared to that of Col-0, one aliphatic GSL compound, 4OHB exhibited reduced amount in both *tpl* and *tpr1/tpl/tpr4* mutants compared to that of Col-0. **C)** Quantification of four different indolic GSL compounds between Col-0, *tpl*, and *tpl/tpr1/tpr4* mutant seedlings. Four indolic GSL compounds displayed a complicated patterns when compared between Col-0, *tpl* and *tpl/tpr1/tpr4* mutants. **D)** Quantification of total aliphatic GSLs, total indolic GSLs and total GSLs (combining aliphatic and indolic GSLs) compounds between Col-0 and the *hda19-3,* a knock-out mutant of *HDA19*. Amount of total aliphatic GSL compounds were substantially higher in the *hda19-3* mutant compared ot that of Col-0. However, amount of total indolic GSL compounds was not signifiantly different between Col-0 and the *hda19-3* mutant. Amounts of total GSLs were calculated by combining amounts of aliphatic and indolic GSL compounds. Alip: aliphatic, Ind: indolic. **E)∼F)** Quantification of four aliphatic GSLs (**E**) and four indolic GSLs (**F**) between Col-0 and *hda19-3* mutant. Abbreviations: 4OHB, 4-Hydroxybutyl; 4MSOB, 4-Methylsulfinylbutyl; 5MSOP, 5-Methylsulfinylpentyl; 4MTB, 4-Methylthiobutyl; I3M, Indol-3-ylmethyl; 4OHI3M, 4-Hydroxyindol-3-ylmethyl; 4MOI3M, 4-Methoxyindol-3-ylmethyl; 1MOI3M, 1-Methoxyindol-3-ylmethyl. **G)** Result of ChIP-PCR analysis using anti-GFP antibody (ab290, Abcam, UK) on aliphatic GSL pathway genes, *MYB29, CYP79F1, CYP83A1*, and *FMOGS-OX1* between *bzr1-ko* mutant and pBZR1::BZR1-CFP (W2C) transgenic line. Genomic structure of individual gene and relative location of PCR amplicons used for ChIP-qPCR analysis (indicated with PX, X is arabic number) were shown above each bar graph. Eriched level of individual PCR amplicon in the *bzrk-ko* mutant was converted to 1, and then relative enriched level of individual PCR amplicon of the pBZR1::BZR1-CFP (W2C) transgenic line were indicated in each graph. Horizontal red dashed line indicates the level of normalized level of *bzr1-ko* mutant. **H)** Result of ChIP-PCR analysis using anti-TPL antibody (AS16 3207, Agrisera, Sweden) on *MYB29* between *tpl* mutant and Col-0 wild type. Genomic structure of individual gene and relative location of PCR amplicons used for ChIP-qPCR analysis (indicated with PX, X is arabic number) were shown above each bar graph. Eriched level of individual PCR amplicon in the *tpl* mutant was converted to 1, and then relative enriched level of individual PCR amplicon of the Col-0 wild type were indicated in each graph. Horizontal red dashed line indicates the level of normalized level of *tpl* mutant. **I)** Result of RT-pPCR analysis on *MYB29* between Col-0 and *tpl* mutant. **J)** Result of ChIP-PCR analysis using anti-H3ac antibody (06-599, Merck, Germany) on *MYB29* between Col-0 and *hda19-3* mutant. Genomic structure of individual gene and relative location of PCR amplicons used for ChIP-qPCR analysis (indicated with PX, X is arabic number) were shown above each bar graph. Eriched level of individual PCR amplicon in the *tpl* mutant was converted to 1, and then relative enriched level of individual PCR amplicon of the Col-0 wild type were indicated in each graph. Horizontal red dashed line indicates the level of normalized level of *tpl* mutant. **K)** Result of RT-qPCR analysis on *MYB29* between Col-0 and *hda19-3* mutant. **A)∼K)** Two-week-old seedlings grown under long day (16h light: 8h dark) at 22℃ were harvested at ZT4 and subjected for molecular experiments like HPLC, ChIP-qPCR, and qRT-PCR analyses. One-way ANOVA with Tukey’s post-hoc test was applied to calculate the statistical significance, and significant difference was indicated in the figures by different letters (p < 0.05). Data were presented as means ± standard deviation (SD) of three biological replicates.

Among the four detected aliphatic GSL compounds, 4MTB exhibited the most pronounced increase in both mutants relative to Col-0 and accounted largely for the elevated total aliphatic GSL levels (Fig. 2B). In addition, two aliphatic GSLs, 4MSOB and 5MSOP, were also increased in both *tpl* and *tpl;tpr1;tpr4* mutants, whereas 4OHB was consistently reduced compared with Col-0. As a result, the total aliphatic GSL content was significantly higher in both mutants than in the wild type. In contrast, total indolic GSL levels were not significantly altered in either *tpl* or *tpl;tpr1;tpr4* mutants; however, individual indolic GSL compounds displayed distinct and variable changes relative to Col-0 (Fig. 2C). Collectively, these results indicate that loss of *TPL* affects aliphatic and indolic GSL accumulation, exerting a strong effect on aliphatic GSL levels while causing more variable changes in individual indolic GSL compounds.

Next, we quantified the levels of aliphatic and indolic GSLs in Col-0 and *hda19-3*, a loss-of-function mutant of the histone deacetylase HDA19. The total aliphatic GSL content was significantly higher in *hda19-3* mutant than in Col-0, whereas total indolic GSL levels were not significantly altered (Fig. 2D-2F). These results suggest a role for HDA19 in the negative regulation of aliphatic GSL accumulation. Taken together with our analyses of *bzr1-1D* and *tpl* mutants, these data suggest that BZR1 acts in concert with the TPL–HDA19 co-repressor complex to negatively modulate GSL biosynthesis, particularly the aliphatic GSL pathway.

Previous studies have shown that BZR1 and BES1 directly interact with a group of basic helix-loop-helix (bHLH) transcription factors, BES1-INTERACTING MYC-LIKE 1 (BIM1), BIM2, and BIM3, to regulate a subset of BR-responsive genes (Yin et al., 2005). To assess whether BIM proteins are involved in the regulation of GSLs metabolism, we measured endogenous aliphatic and indolic GSL levels in a *bim1;bim2;bim3* triple mutant (hereafter *bim1;2;3*), in which functional redundancy among BIMs is eliminated. No significant differences in either aliphatic or indolic GSL levels were observed in *bim1;2;3* compared with Col-0 (Supplementary Fig. S2). We further examined the involvement of additional BR-responsive bHLH transcription factors by analyzing GSL levels in the *bee1;bee2;bee3* triple mutant (*bee1;2;3*) (Friedrichsen et al., 2002). Similar to *bim1;2;3*, the *bee1;2;3* mutant did not exhibit significant changes in aliphatic or indolic GSL content relative to Col-0 (Supplementary Fig. S2). These results indicate that BIM and BEE family bHLH transcription factors are unlikely to play major roles in BR-mediated regulation of GSL biosynthesis in *Arabidopsis*.

### BZR1-TPL(R)-HDA19 module directly binds to the promoter region of *MYB29* controlling aliphatic GSL biosynthesis

A recent study demonstrated that BZR1 directly binds to the promoter regions of *MYB29* for the regulation of aliphatic GSLs biosynthesis (Wang et al. 2023). In a consistency with this report, ChIP-followed qPCR (ChIP-qPCR) analysis using the *pBZR1::BZR1-CFP* transgenic plant confirmed that *MYB29* was substantially bound by BZR1-GFP (Fig. 2G). However, we could not detect enrichment of BZR1-CFP on other aliphatic GSL pathway genes like *CYP79F1*, *CYP83A1*, and *FMOGS-OX1*. Next, because BZR1-CFP was shown to significantly bind to *MYB29*, we examined whether TPL also directly bind to the promoter region of *MYB29* by conducting ChIP-qPCR analysis using anti-TPL antibody. As a result, TPL was shown to significantly bind to the promoter region of *MYB29*, in a similar region of BZR1-CFP (Fig. 2H). In addition, we also examined whether TPR1, a TPL paralog directly binds to the genomic region of *MYB29* by conducted another round of ChIP-qPCR analysis using *pTPR1::TPR1-HA/tpr1-1* transgenic line. Evident enrichment of TPR1-HA was detected on the promoter of *MYB29* compared to input DNA samples (Supplementary Fig. S3A), suggesting that a TPL paralog, TPR1 (and possibly TPR4) participate in the transcriptional regulation of *MYB29*, even we could not observe synergistic effect of *tpl;tpr1;tpr4* compared to *tpl* mutant in terms of total amounts of aliphatic GSLs (Fig. 2A). In a consistency with direct binding of BZR1 and TPL (and TPR1), transcript level of *MYB29* was upregulated in the *tpl* (Fig. 2I) and *tpl;tpr1;tpr4* mutant in comparison to those of Col-0 (Supplementary Fig. S3B).

### BZR1-TPL(R)-HDA19 module suppresses *MYB29* via histone deacetylation

As a histone deacetylase, HDA19 catalyzes the removal of acetyl groups from histone H3, thereby contributing to transcriptional repression (Jiang et al., 2020). Given the increased accumulation of aliphatic GSLs observed in the *hda19-3* mutant relative to Col-0, we hypothesized that HDA19 epigenetically suppresses the expression of *MYB29* by removing histone acetylation marks from its chromatin. To test this hypothesis, we performed chromatin immunoprecipitation followed by quantitative PCR (ChIP-qPCR) using an anti-H3ac antibody to assess H3 acetylation levels at the *MYB29* locus in Col-0 and *hda19-3*. A substantial enrichment of H3ac at *MYB29* was detected in *hda19-3* compared with Col-0 (Fig. 2J). Consistent with this increased histone acetylation, *MYB29* transcript levels were significantly elevated in *hda19-3* relative to Col-0 (Fig. 2K). These results indicate that the BZR1-TPL(R)-HDA19 regulatory module epigenetically represses *MYB29* expression by reducing H3 acetylation at BZR1 chromatin, thereby negatively regulating aliphatic GSLs biosynthesis in *Arabidopsis*.

### UBP12 and UBP13, deubiquitinases of BZR1 play a negative role in GSLs biosynthesis in Arabidopsis

The deubiquitinating enzymes, UBP12 and UBP13 function as positive regulators of BR signaling by removing ubiquitin from key BR signaling components, including BES1 and BRI1 (Luo et al., 2022, Park et al., 2022). In addition, UBP12 and UBP13 were previously identified as BZR1-associated proteins through tandem affinity purification in *Arabidopsis* (Wang et al., 2013a). These findings raised the possibility that UBP12/13-mediated de-ubiquitination of BZR1 may contribute to the regulation of GSLs biosynthesis. To test this hypothesis, we quantified GSL levels in Col-0 and the *ubp12w;ubp13* double mutant, which is defective in both *UBP12* and *UBP13*. HPLC analysis revealed that the total GSL content, comprising both aliphatic and indolic GSLs, was significantly increased in *ubp12w;ubp13* compared with Col-0 (Fig. 3A). Among aliphatic GSLs, three compounds (4OHB, 4MSOB, and 4MTB) accumulated to higher levels in the *ubp12w;ubp13* mutant than in the wild type, whereas 5MSOP, the least abundant aliphatic GSL detected, did not differ significantly between genotypes (Fig. 3B). For indolic GSLs, two major compounds, I3M and 1MOI3M, were present at higher levels in *ubp12w;ubp13* than in Col-0, while 4OHI3M and 4MOI3M showed no significant changes (Fig. 3C). Consistent with these metabolic changes, RT-qPCR analysis revealed that several aliphatic GSL biosynthetic genes, including *MYB28*, *MYB29*, *CYP83A1*, *BCAT4*, and *MAM1*, were significantly upregulated in the *ubp12w;ubp13* mutant relative to Col-0 (Fig. 3D). Together, these results suggest that UBP12 and UBP13 negatively influence GSL biosynthesis, potentially by stabilizing BZR1 and thereby enhancing BZR1-mediated repression of GSL biosynthetic gene expression in *Arabidopsis*.

**Figure 3.**
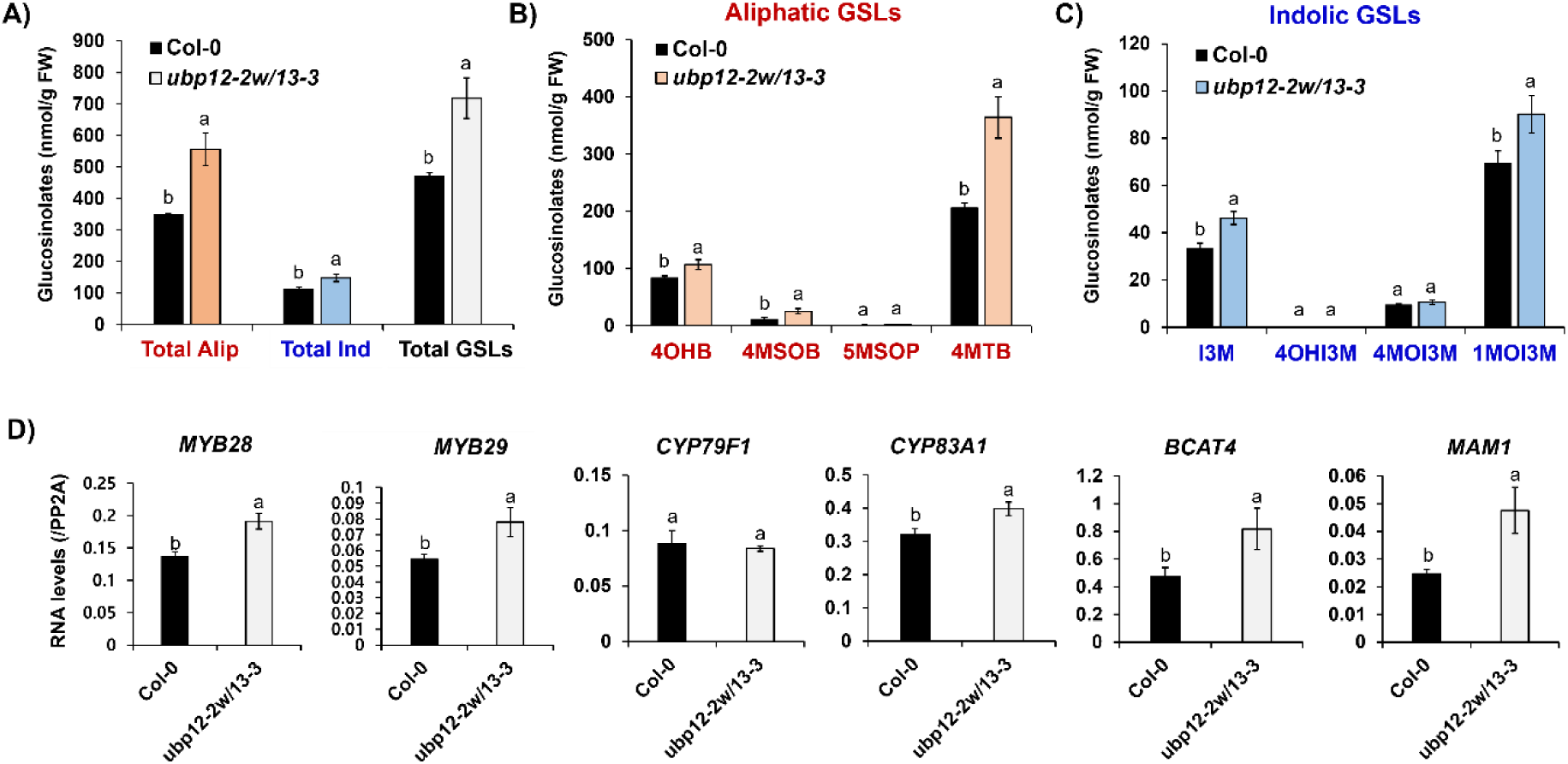
UBP12 and UBP13, two deubiquitinases of BZR1 negatively affect GSLs biosynthesis. **A)** Quantification of aliphatic and indolic GSL compounds between Col-0 and the *ubp12-2w;ubp13-3,* a double mutant of *UBP12* and *UBP13*. Amounts of total GSLs were calculated by combining amounts of aliphatic and indolic GSL compounds. Alip: aliphatic, Ind: indolic. **B)** Quantification of aliphatic GSL compounds between Col-0 and the *ubp12-2w;ubp13-3* mutant. **C)** Quantification of indolic GSL compounds between Col-0 and the *ubp12-2w;ubp13-3* mutant. **D)** Result of RT-qPCR analysis on six aliphatic GSL pathway genes between Col-0 and *ubp12-2w;ubp13-3* mutant. **A)∼D)** Two-week-old seedlings grown under long day (16h light: 8h dark) at 22℃ were harvested at ZT4 and subjected for HPLC and RT-qPCR analyses. One-way ANOVA with Tukey’s post-hoc test was applied to calculate the statistical significance, and significant difference was indicated in the figures by different letters (p < 0.05). Data were presented as means ± standard deviation (SD) of three biological replicates.

### Abscisic acid (ABA) signaling acts to promote GSLs biosynthesis

Given that BR signaling negatively regulates GSLs biosynthesis, we next investigated which plant hormones might antagonize BR-mediated suppression of GSL accumulation. To address this, we examined the effects of five plant hormones, brassinosteroid (BR; 24-epibrassinolide, EBR), abscisic acid (ABA), gibberellic acid (GA), methyl jasmonate (MeJA), and auxin (1-naphthaleneacetic acid, NAA) on the accumulation of aliphatic and indolic GSLs. For aliphatic GSLs, ABA and MeJA treatments resulted in a significant increase in aliphatic GSL levels, whereas EBR, GA, and NAA treatments led to reduced accumulation compared with Col-0 controls (Fig. 4A). Similarly, for indolic GSLs, ABA- and MeJA-treated samples exhibited significantly higher levels than those treated with other hormones (Fig. 4B). In contrast, NAA treatment caused a pronounced reduction in indolic GSL accumulation, while EBR and GA treatments did not result in significant changes relative to Col-0. Consequently, these five plant hormones differentially affected total endogenous GSL levels, calculated as the sum of aliphatic and indolic GSLs (Fig. 4C). Notably, ABA exerted a positive effect on both aliphatic and indolic GSL accumulation. This observation is consistent with previous reports suggesting that ABA antagonizes BR-mediated regulation of plant growth and development in *Arabidopsis* (Wang et al., 2020, Wang et al., 2018, Wu et al., 2020).

**Figure 4.**
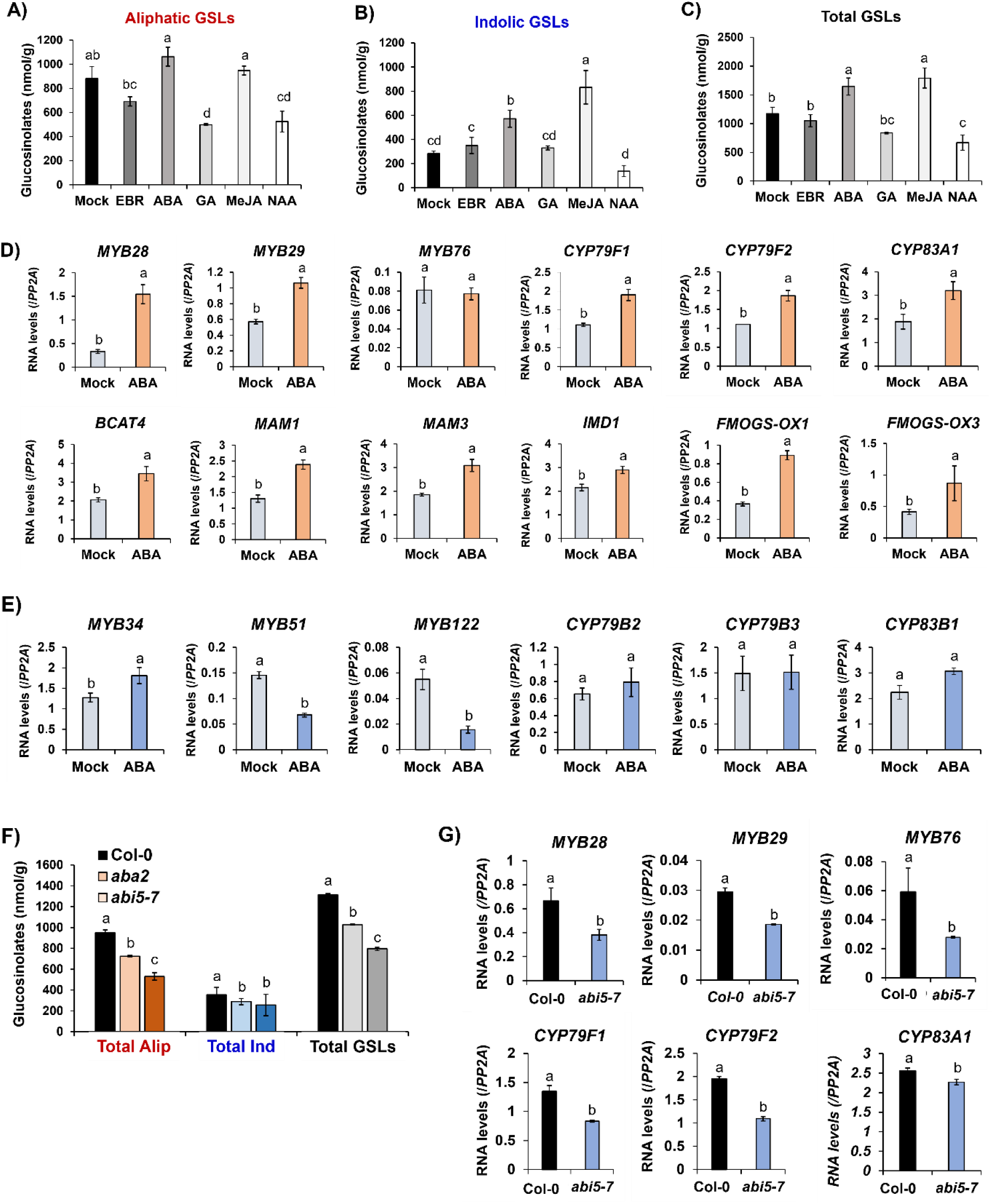
ABI5-mediated ABA signaling plays a positive role in the regulation of aliphatic GSL pathway genes. **A)** Quantification of aliphatic GSL compounds between control Col-0 Col-0 (Mock), EBR (5uM), ABA (5uM), GA (10uM), MeJA (50uM), and NAA (5uM)-treated samples. **B)** Quantification of indolic GSL compounds between control Col-0 (Mock), EBR (5uM), ABA (5uM), GA (10uM), MeJA (50uM), and NAA (5uM)-treated samples. **C)** Comparison of total GSL compounds (combining aliphatic and indolic GSL compounds) between Col-0 (Mock), EBR (5uM), ABA (5uM), GA (10uM), MeJA (50uM), and NAA (5uM)-treated samples. **A)∼C)** One-week-old seedlings grown under long day (16h light: 8h dark) at 22℃ were transferred to each hormone-containing MS media and grown further 3 days. Mock or hormone-treated seedlings were harvested at ZT4 and subjected for HPLC and RT-qPCR analyses. **D)** Result of qRT-PCR analysis of 16 aliphatic GSL pathway genes between Mock (untreated) and ABA-treated samples. **E)** Result of qRT-PCR analysis of six indolic GSL pathway genes between Mock (untreated) and ABA-treated samples. **F)** Quantification of aliphatic and indolic GSL compounds between Col-0, *aba2* (a knockout mutant of ABA biosynthetic gene, *ABA2*), and the *abi5-7* (a knock-out mutant of *ABI5*). Amounts of total GSLs were calculated by combining amounts of aliphatic and indolic GSL compounds. Two-week-old seedlings grown under long day (16h light: 8h dark) at 22℃ were harvested at ZT4 and subjected for HPLC analysis (n =3). One-way ANOVA with Tukey’s post-hoc test was applied to calculate the statistical significance, and significant difference was indicated in the figures by different letters (p < 0.05). Alip: aliphatic, Ind: indolic. **G)** Result of RT-qPCR analysis on six aliphatic GSL pathway genes between Col-0 and *abi5-7* mutant. With the exception of *MYB28*, five tested aliphatic GSL pathway genes (*MYB29, MYB76, CYP79F1, CYP79F2*, and *CYP83A1*) showed reduced transcript levels in the *abi5-7* mutant Compared to Col-0. Two-week-old seedlings grown under long day (16h light: 8h dark) at 22℃ were harvested at ZT4 and subjected for RT-qPCR analysis (n =3). One-way ANOVA with Tukey’s post-hoc test was applied to calculate the statistical significance, and significant difference was indicated in the figures by different letters (p < 0.05).Alip: aliphatic, Ind: indolic. **A)∼G)** One-way ANOVA with Tukey’s post-hoc test was applied to calculate the statistical significance, and significant difference was indicated in the figures by different letters (p < 0.05). Data were presented as means ± standard deviation (SD) of three biological replicates.

To determine whether ABA promotes GSL biosynthesis through transcriptional activation of GSL pathway genes, we performed RT-qPCR analysis comparing ABA-treated and mock-treated Col-0 seedlings. Among the 16 examined aliphatic GSL biosynthetic genes, a substantial proportion (10 out of 16 genes; 63%) were significantly upregulated following ABA treatment, whereas the remaining six genes showed no significant changes (Fig. 4D). These results indicate that ABA positively regulates the expression of a large subset of aliphatic GSL biosynthetic genes. In contrast, expression patterns of six indolic GSL pathway genes were more variable. For example, *MYB34* was upregulated in ABA-treated samples, whereas *MYB51* and *MYB122* were downregulated relative to mock-treated controls (Fig. 4E). Other indolic GSL biosynthetic genes, including *CYP79B2*, *CYP79B3*, and *CYP83B1*, did not exhibit significant expression changes.

ABI5, a key transcription factor in ABA signaling, has been reported as a major downstream target of BR signaling (Skubacz et al., 2016). To further investigate the role of ABA signaling in GSL biosynthesis, we compared GSL levels among Col-0, *aba2* (a loss-of-function mutant in the ABA biosynthetic gene, *ABA2*), and *abi5-7* (a loss-of-function mutant of *ABI5*). Aliphatic GSL levels were substantially reduced in both *aba2* and *abi5-7* mutants compared with Col-0 (Fig. 4F). Similarly, total indolic GSL levels were also lower in these mutants than in the wild type. Consistent with these metabolic changes, RT-qPCR analysis revealed that several aliphatic GSL biosynthetic genes, including *MYB29*, *MYB76*, *CYP79F1*, *CYP79F2*, and *CYP83A1*, were expressed at significantly lower levels in *abi5-7* than in Col-0 (Fig. 4G). Taken together, these results demonstrate that ABA signaling positively regulates GSL biosynthesis and that ABI5 functions as a key positive regulator of both aliphatic and indolic GSL production in *Arabidopsis*.

### ABI5 directly binds to the promoter region of BR signaling components, *BZR1* and *BES1* and *UBP12* and *UBP13* to inhibit BR-mediated repression of GSLs biosynthesis in Arabidopsis

Given that ABI5 functions antagonistically to BZR1/BES1-mediated repression of GSLs biosynthesis, we first examined whether ABI5 physically interacts with BZR1 and/or BES1 to interfere with their functions. To test this possibility, we performed yeast two-hybrid (Y2H) assays. However, no direct interaction between ABI5 and either BZR1 or BES1 was detected (Supplementary Fig. S4). Because ABI5 acts as a transcription factor in ABA signaling, we next explored whether ABI5 directly regulates GSL biosynthetic genes at the transcriptional level. To this end, we analyzed a publicly available ABI5 ChIP-seq dataset generated using an anti-ABI5 antibody (GEO accession number: GSM1466341) to identify genome-wide ABI5 binding sites. No significant ABI5 enrichment was observed at any aliphatic GSL biosynthetic gene loci (Supplementary Fig. S5A). In contrast, for indolic GSL pathway genes, prominent ABI5 binding peaks were detected near the transcription start sites of *MYB34* and *GSTF10* (Supplementary Fig. S5B). These observations suggest that ABI5 may directly regulate a subset of indolic GSL biosynthetic genes, potentially contributing to ABI5-mediated activation of indolic GSL biosynthesis. Further investigation will be required to elucidate the functional significance of these interactions.

Because ABI5 binding was not detected at aliphatic GSL pathway genes, we next examined whether ABI5 instead regulates components of the BR signaling pathway. Notably, strong ABI5 enrichment was detected at the genomic regions of key BR signaling genes, including *BES1*, *BZR1*, *UBP12*, and *UBP13*, in the ChIP-seq dataset (Fig. 5A–B). To validate these observations, we performed ChIP–qPCR analysis using an anti-ABI5 antibody in Col-0 and the *abi5-7* mutant under both mock and ABA-treated conditions. Under mock conditions, no significant ABI5 enrichment was detected at the *BES1* or *BZR1* loci (Fig. 5C). In contrast, under ABA treatment, ABI5 showed substantial enrichment at the genomic regions of *BES1*, *BZR1*, *UBP12*, and *UBP13* (Fig. 5D). To determine whether ABI5 binding affects transcription of these BR signaling genes, we performed RT–qPCR analysis comparing Col-0 and *abi5-7*. Transcript levels of *BES1*, *BZR1*, *UBP12*, and *UBP13* were significantly elevated in the *abi5-7* mutant relative to Col-0 (Fig. 5E). Taken together, these results indicate that ABI5 negatively regulates the expression of multiple BR signaling components, including *BZR1*, *BES1*, *UBP12*, and *UBP13*, thereby providing a mechanistic basis for ABA-mediated antagonism of BR signaling in the regulation of GSL biosynthesis.

**Figure 5.**
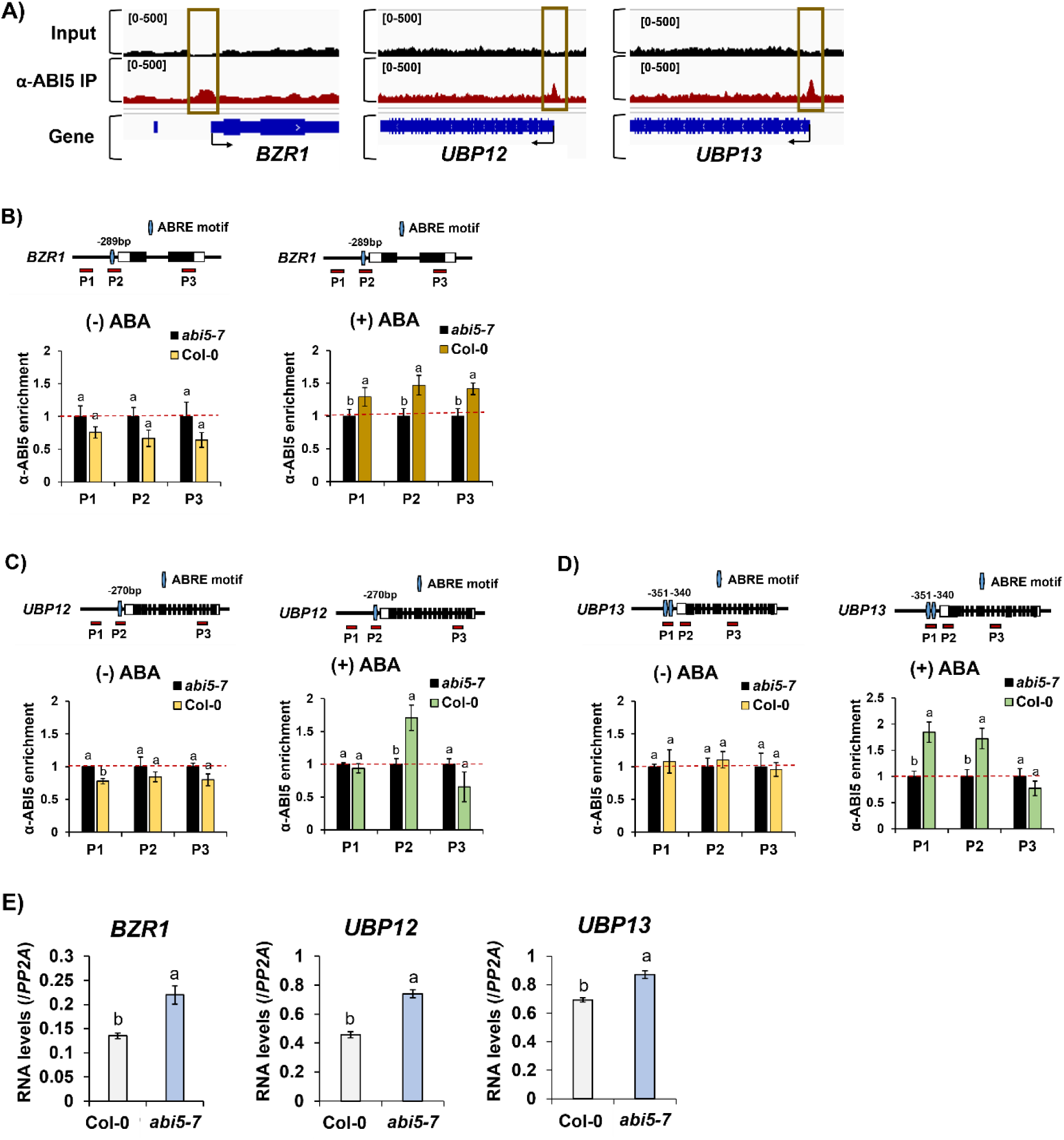
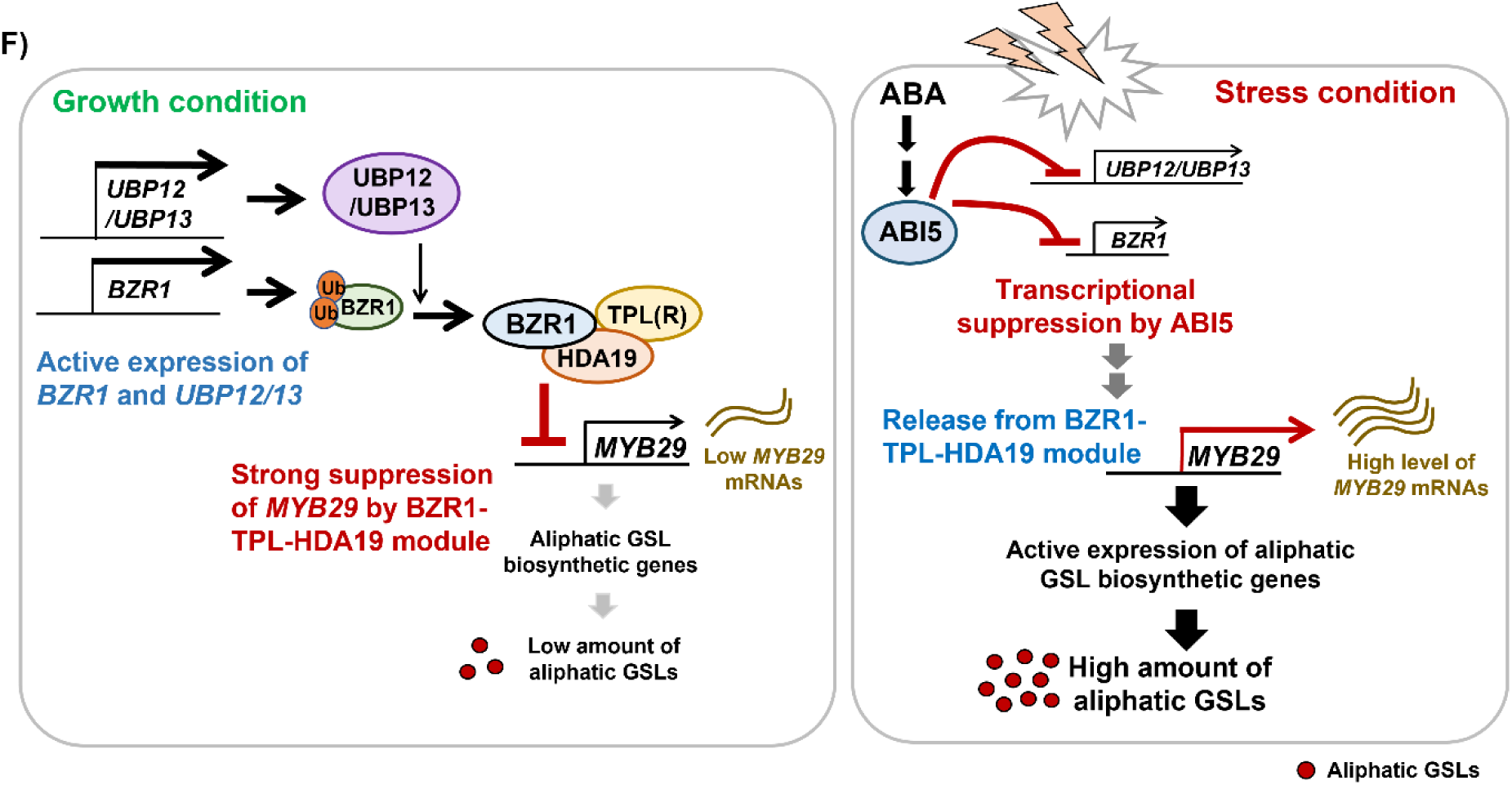
ABI5 directly regulate expression of two BR signaling factors, *BZR1* and *BES1* as well as deubiquitinase genes, *UBP12* and *UBP13*. **A)** Analysis of a public ChIP-seq dataset using α-ABI5 on key BR signaling factor, *BZR1*. The IGV genome browser view illustrates significant enrichment of ABI5 binding on the proximal promoter region of *BZR1*. A green box highlights the regions with significantly enriched ABI5 peaks. Input DNA and α-ABI5-immunoprecipitated DNA are shown in black and red tracks, respectively. The range of binding intensity on the Y-axis is indicated by brackets at the upper left corner of each track. **B)** Analysis of a public ChIP-seq dataset using α-ABI5 on two deubiquitinase genes, *UBP12* and *UBP13* which encodes deubiquitinases for BZR1, thus reinforcing BR signaling by stabilizing BZR1. The IGV genomic browser view illustrates significant enrichment of ABI5 binding on the proximal promoter region of *UBP12* and *UBP13*. A green box highlights the regions with significantly enriched ABI5 peaks. Input DNA and α-ABI5-immunoprecipitated DNA are shown in black and red tracks, respectively. The range of binding intensity on the Y-axis is indicated by brackets at the upper left corner of each track. **C)** Result of ChIP-qPCR analysis using α-ABI5 on key BR signaling gene, *BZR1* as well as two deubiquitinase genes, *UBP12* and *UBP13* under no-ABA (-ABA) condition. D) Result of ChIP-qPCR analysis using α-ABI5 on key BR signaling gene, *BZR1* as well as two deubiquitinase genes, *UBP12* and *UBP13* under ABA (5uM)-treated condition. C)∼D) Genomic structure of individual gene and relative location of PCR amplicons used for ChIP-qPCR analysis (indicated by red line with labeling of PX, X is arabic number) were shown above each bar graph. Horizontal red dashed line indicates the level of normalized enrichment of level of α-ABI5-immunoprecipitated DNA in *abi5-7* mutant. **E)** Result of qRT-PCR analysis on *BZR1, UBP12,* and *UBP13* genes between Col-0 and the *abi5-7* mutant. Transcript level of each gene was normalized by the comparison to level of reference gene, *PP2A* (AT1G13320). **C)-E)** One-way ANOVA with Tukey’s post-hoc test was applied to calculate the statistical significance, and significant difference was indicated in the figures by different letters (p < 0.05). Data were presented as means ± standard deviation (SD) of three biological replicates. **F)** A schematic model underlying the crosstalk between BR and ABA signaling in the regulation of GSLs biosynthesis. In normal (unstressed) growth condition (left panel), BZR1 is actively involved in the promotion of plant growth and development. During the growth condition, UBP12/UBP13 functions to actively de-ubiquitinate BZR1, thus stabilizing its protein stability. Stabilized BZR1 directly binds to the genomic region of *MYB29*, thus resulting in the suppression of aliphatic GSLs biosynthesis. As a result, plants have low amount of aliphatic GSLs in normal (growth) condition. Meanwhile, upon stressed (ABA abundant) condition (right panel), plants ceases growth and development, instead promoting defensive processes including increasing amounts of aliphatic GSLs. ABA-stabilized ABI5 actively bind and suppress transcription of *BZR1* as well as *UBP12* and *UBP13,* two deubiquitinase genes responsible for stabilizing BZR1. As a result, de-repressed *MYB29* reinforces production of aliphatic GSLs. This ABI5 action results in the high accumulation of aliphatic GSLs in a stressed (ABA) condition.

Based on these results, we came up with a schematic model for the regulatory mechanism underlying the ABI5-mediated enhancement of GSLs biosynthesis via transcriptional suppression of BR signaling factors in Arabidopsis (Fig. 6I). In unstressed (growth) condition (left panel), BZR1 are actively involved in the promotion of plant growth and development. During the growth condition, BZR1 forms suppressive module with TPL and HDA19. Biosynthesis of aliphatic GSLs is suppressed by BZR1-TPL/HDA19 module which mediates repression of a MYB TF, *MYB29*, thus resulting in the low amount of aliphatic GSLs (left panel). Meanwhile, upon some stressed (ABA abundant) condition (right panel), plants ceases growth and development, instead promoting defensive processes including increasing amounts of aliphatic GSLs. In this process, ABI5 actively suppresses transcription of BR signaling factor, *BZR1* and two deubiquitinase genes (e.g. *UBP12* and *UBP13*), which play a negative role in biosynthesis of aliphatic GSLs in *Arabidopsis*. Resultantly, de-repressed *MYB29* promotes expression of downstream biosynthetic genes for aliphatic GSLs, thus resulting in the accumulation of aliphatic GSLs. Resultantly, this ABI5 action results in the high accumulation of aliphatic GSLs in a stressed (ABA) condition (right panel).

## DISCUSSION

BZR1 and BES1 are key transcription factors in the brassinosteroid (BR) signaling pathway and regulate the expression of thousands of genes in *Arabidopsis* (Kim and Wang, 2010). Although the roles of BZR1 in BR-mediated plant growth and development have been well documented, the molecular mechanisms by which BZR1 regulates secondary metabolism remain poorly understood. In this study, we demonstrates that BZR1 functions as negative regulators of GSLs biosynthesis through their association with TPL/TPR corepressors and the histone deacetylase, HDA19. Notably, BZR1 plays a predominant role in repressing aliphatic GSLs biosynthesis by epigenetically suppressing *MYB29* expression. Furthermore, we show that the ABA signaling factor ABI5 directly represses the expression of BR signaling genes, including *BZR1*, *UBP12*, and *UBP13*, thereby positively modulating the expression of GSL biosynthetic genes in *Arabidopsis*.

Regarding the negative activity of BZR1 in the regulation of GSL pathway genes, EAR motifs within BZR1 exert as a transcription repression domain by recruiting the Groucho/TLE-like transcriptional corepressor, TPL (Oh et al., 2014). In our study, we found that *tpl,* a knockout mutant of *TPL* exhibited a significant increase of total amounts of GSLs, indicating that TPL is involved in the BZR1-mediated suppression of GSL pathway genes in *Arabidopsis*. In addition, because TPL was previously reported to share a functional redundancy with its paralogs like TPR1 and TPR4, we further compared endogenous amounts of GSLs between Col-0 and *tpl;tpr1;tpr4* triple mutants. we could not observe significant increase of aliphatic GSLs in *tpl;tpr1;tpr4* triple mutant compared to those of the *tpl* single mutant (Fig. 2A-2C). It might indicate that TPL does not share functional redundancy with its paralogs, TPR1 and TPR4 in the regulation of GSLs biosynthesis in *Arabidopsis*.

Although BZR1 gain-of-function resulted in reduced accumulation of both aliphatic and indolic GSLs (Fig. 1D), loss of *TPL* and its paralogs led to a marked increase in aliphatic GSLs without a significant change in total amounts of indolic GSLs (Fig. 2A-2C). This discrepancy suggests that BZR1-mediated repression of aliphatic GSLs biosynthesis largely depends on the TPL/TPR co-repressor module, whereas regulation of indolic GSLs biosynthesis may involve additional layers of control. Notably, individual indolic GSL compounds displayed dynamic and opposing changes in *tpl* and *tpl;tpr1;tpr4* mutants, indicating that loss of TPL and its paralogs might affect the composition of indolic GSLs rather than the overall pool size. Such compensatory redistribution among indolic GSL compounds might mask changes in total amounts of indolic GSLs. Furthermore, it is possible that BZR1 might regulate indolic GSLs biosynthesis through TPL-independent mechanisms, potentially involving alternative co-regulators or indirect transcriptional networks that contribute to maintaining indolic GSLs homeostasis. Together, these observations highlight distinct regulatory modes by which BZR1 controls aliphatic and indolic GSL biosynthesis in *Arabidopsis*.

BZR1 have been shown to utilize the TPL-HDA19 corepressor module to suppress the expression of defense-related pathway genes, including those involved in glucosinolates (GSLs) biosynthesis (Kim et al., 2019, Ryu et al., 2014, Oh et al., 2014). Under normal, non-stress conditions, BZR1 primarily promote plant growth and development by actively regulating cell division and elongation. In contrast, under stress conditions characterized by elevated abscisic acid (ABA) levels, plants suppress growth and instead prioritize defensive responses against environmental stresses such as heat, salinity, and drought (Manghwar et al., 2022, Planas-Riverola et al., 2019). Consistent with this growth–defense trade-off, the biosynthesis of defensive metabolites, including GSLs, is attenuated by the BZR1-TPL-HDA19 repressive module. An important remaining question is whether the BZR1-TPL-HDA19 complex also regulates additional metabolic pathways beyond GSL biosynthesis, particularly in response to distinct types of environmental stress.

GSLs metabolism was dynamically affected by different plant hormones (Fig 4A-4C). Among plant hormones, jasmonates (JAs) plays important roles in plant defense against a diversity of stresses by triggering production of defensive secondary metabolites including GSLs (Kazan, 2015). In particular, three basic helix-loop-helix TFs, MYC2, MYC3 and MYC4 execute a major function in JA-mediated production of GSLs in *Arabidopsis* plant. A previous study demonstrated that these MYC TFs are positively required for expression of many GSLs pathway genes (Schweizer et al., 2013). In addition, MYCs was shown to interact with MYB TFs like MYB28, MYB29, MYB76, MYB34, MYB51, and MYB122, thus forming MYBs-MYCs heterodimer to positively regulate expression of aliphatic and indolic GSL pathway genes. Because JA signaling affect GSL metabolism, we examined whether JA signaling components like MYC2, MYC3, and MYC4 affect expression of BR signaling components like *BZR1, UBP12*, and *UBP13* by conducting RT-qPCR analysis between Col-0, *myc2*, and *myc2;3;4* mutants of *Arabidopsis* plants. As a result, the transcript levels *of BZR1, UBP12,* and *UBP13* did not show significant differences among *Col-0*, *myc2*, and *myc2;3;4* mutants, with *UBP12* and *UBP13* showing a slightly lower expression in the *myc2;3;4* mutant, rather than being elevated (Supplementary Fig. S6A). In addition, analysis of public available RNA-seq dataset between Col-0 and *myc2;3;4* mutant also showed that transcript levels of *BZR1, UBP12*, and *UBP1*3 were not substantially altered between Col-0 and *myc2;3;4* mutant (Supplementary Fig. S6B) (Van Moerkercke et al., 2019). Collectively, these results reflect that JA signaling might impact on GSL biosynthesis in a different manner from the ABI5-mediated promotion of GSLs biosynthesis in *Arabidopsis*.

In conclusion, the crosstalk between ABA and BR signaling pathways allows plants to fine-tune their responses to changing environmental conditions by integrating signals from different hormonal pathways. This integration enables plants to optimize their growth and development while coping with various stresses.

## MATERIALS AND METHODS

### Plant materials and growth conditions

*Arabidopsis thaliana* ecotypes Columbia-0 (Col-0), Enkheim-2 (En-2), mutants, and transgenic plants were grown in the growth room at 22 °C in long day condition (16h day; 8h dark) or short day condition (8h day: 16h dark). White light was supplied by cool-white fluorescent lamps (FHF322SSEX-D, 120 μmol m^−2^ s^−1^; Osram, Republic of Korea). The *abi5-7* mutation that causes a premature stop codon at W^75^ in the *ABI5* gene was previously described (Nambara *et al*., 2002). Higher order mutants, *bim1;2;3* and *bee1;2;3* (Friedrichsen et al., 2002), *myc2;3;4* triple mutant (Schweizer et al., 2013), *tpl;tpr1;tpr4* triple mutant (Zhu et al., 2010), and gain-of-function mutants, *bzr1-1D* (Wang et al., 2002), *bes1-D* (Yin et al., 2002) were previously reported. The *tpl* (SALK_034518), *aba2* (CS156), *hda19-3* (SALK_139445), and *hac1-1* (SALK_082118) were obtained from the Arabidopsis Biological Resource Center (ABRC). Transgenic plants of *pBZR1-BZR1-CFP* (W2C) was previously reported (Oh et al., 2012). *bzr1-ko* mutant is a null mutant containing 90bp-deletion at the first exon region of *BZR1*. The *ubp12-2w* (GABI_742C10), *ubp13-3* (SALK_132368), *ubp12-2w/13-3* (a double mutant) were previously reported (Cui *et al*., 2013).

### Quantification of GSLs compounds

Seedlings grown for 14 days in long day condition were sampled at ZT4 for extraction of GSLs. Quantification of GSLs compounds was performed as previously described (Nugroho et al., 2020). 0.5mg/ml of sinigrin (Sigma-Aldrich, USA) was used as an internal standard compound. Each desulfo-GSL was analyzed with high-performance liquid chromatography (Vanquish Core HPLC system, Thermo scientific, USA). Desulfo-GSLs were separated in a 0.5mL min^−1^ flow rate on a C18 reverse phase column (Zorbax XDB-C18, 4.6 x 250mm, 5μm particle size, Agilent, USA) with a water and acetonitrile gradient system. Individual peaks were identified using standard compounds (Phytoplan, Germany) (Supplementary Table S1). Amount of sinigrin was used for relative quantification (Brown et al., 2003). The samples were analyzed independently with three replicates and presented in nmol g^−1^ on a fresh weight (FW) basis.

### Analysis of GSLs by LC-DAD-ESI-MS

Contents of desulfo-GSLs were analyzed as previously described (Nugroho et al., 2020). In brief, Accela ultra-high-performance liquid chromatography system (Thermo scientific, USA) combined with ion trap mass spectrometer (LTQ Velos Pro, Thermo scientific, USA) were used. The samples (25μL) were separated in C18 reverse phase column (Zorbax XDB-C18, 4.6 x 250mm, 5μm particle size, Agilent, USA) and determined in negative ion mode ([M-H]-). Mass-spectrometry (MS) operation was performed as described below: capillary temperature (275 °C), capillary voltage (5kV), source heater temperature (250 °C), sheath gas flow (35 arb), auxiliary gas flow (5arb), and spectra scanning range (m.z 100∼1500). Molecular mass ion of desulfo-4OHB was identified based on the previous report (DS-GATN m/z 310, 356, 621) (Kusznierewicz et al., 2013, Brown et al., 2003, Petersen et al., 2002).

### Quantitative RT-PCR analysis

Two-week-old seedlings grown in either long day or short day light condition were harvested and immediately frozen in liquid nitrogen. Extraction of total RNAs were performed using RNeasy Plant mini kit (QIAGEN, Germany). DNase I (New England Biolabs, USA) were treated to get rid of contaminated genomic DNAs. Total 5μg of purified RNAs were used for complementary DNA synthesis using EasyScript reverse transcriptase enzyme (TransGen Biotech, China). Quantitative RT-PCR analysis was performed using Sol^TM^ 2x Real-Time PCR Smart mix (SolGent Co., Republic of Korea) in a LineGene 9600 Plus Real-Time PCR system (BioER, China) with the following condition: total 45 cycles of denaturation at 95℃ for 15sec, annealing at 60℃ for 25sec, and extension at 72℃ for 35sec. *PP2A* (AT1G13320) was used as an internal control (Czechowski et al., 2005). The quantification was calculated with the delta Ct method (Livak and Schmittgen, 2001). Information used in the qRT-PCR analysis was listed in the Supplementary Table S2.

### Chromatin immunoprecipitation (ChIP) followed by qPCR (ChIP-qPCR) analysis

Two weeks old whole seedlings were collected and cross-linked in a formaldehyde-containing buffer (10mM Tris-HCl, 0.1 mM EDTA, 0.4 M sucrose, ChIP assay was performed as previously described (Kim and Sung, 2017). Antibody against HA epitope (ab9110, Abcam, UK), ABI5 (ab98831, Abcam, UK), TPL (AS16 3207, Agrisera, Sweden), H3ac histone mark (06-599, Merck Millipore, Germany), and GFP epitope (ab290, Abcam, UK) were purchased and used in this study. Non-transgenic Col-0 or knock-out mutant treated with same antibody was used as a control. Quantitative PCR analysis was performed using BioFACT^TM^ 2XReal-TimePCR Master Mix (BioFACT Co., South Korea) in a LineGene 9600 Plus Real-Time PCR system (Hangzhou BioER Technology Co., China) with the following condition: total 45 cycles of denaturation at 95℃ for 15sec, annealing at 55℃ for 25sec, and extension at 72℃ for 35sec. Amplicon spanning coding sequence of *PP2A* (AT1G13320) was used as an internal control for a relative enrichment quantification. Primer sequences in the ChIP-qPCR analysis was shown in the Supplementary Table S2.

### NGS dataset analysis

Public available ChIP-seq data were downloaded from EBI-EMBL website. The raw FASTQ files were trimmed and quality-filtered. Filtered reads were aligned to the Arabidopsis thaliana TAIR10 reference genome by using Bowtie2 (Langmead and Salzberg, 2012). Aligned reads were converted to digital counts using featureCounts (Liao et al., 2014). Aligned reads were also converted into bigwig files for visualization using IGV genome browser (Thorvaldsdottir et al., 2013). Peak callings were performed by using MACS2 (Zhang et al., 2008). Detailed information on ChIP-seq data used in this study is described in Supplementary Table S3.

### Statistical analysis

All statistical analyses in this study were executed using SAS software (version 9.4; SAS Institute Inc., Cary, NC, USA). Statistical differences were evaluated by one-way analysis of variance (ANOVA) with Tukey’s post-hoc test (*P* < 0.05). Significant differences were denoted by different alphabetical letters. Data were presented as means ± standard deviation (SD) of three biological replicates.

## Acknowledgements

We thanks to Dr. Hojin Ryu at Chungbuk National University for providing the mutant seeds of *bzr1-1D*, and Dr. Eunkyoo Oh at Korea University for *pBZR1-BZR1-CFP* (w2c) seeds. We also thanks to Prof. Yuelin Zhang at the UBC Vancouver for the *pTPR1:TPR1-HA/tpr1-1* transgenic line and *tpl;tpr4;tpr1* triple mutant. This study was supported by the Korea Forestry Promotion Institute (RS-2025-02213366) and by the National Research Foundation of Korea (grant No. RS-2025-16065991) to D.-H. K.

## Author contributions

DC prepared all plant materials and performed genetic and molecular experiments including HPLC analysis; DC and D-HK planned the experiments and conducted bioinformatics analysis; DC and D-HK analyzed the data and wrote the manuscript. All authors have reviewed and approved the final manuscript for submission.

## Data availability statement

The authors declare that all data supporting the findings of this study are available within the manuscript. Supporting information is available from the corresponding author upon request.

## Supplementary Information

**Supplementary Figure S1.**
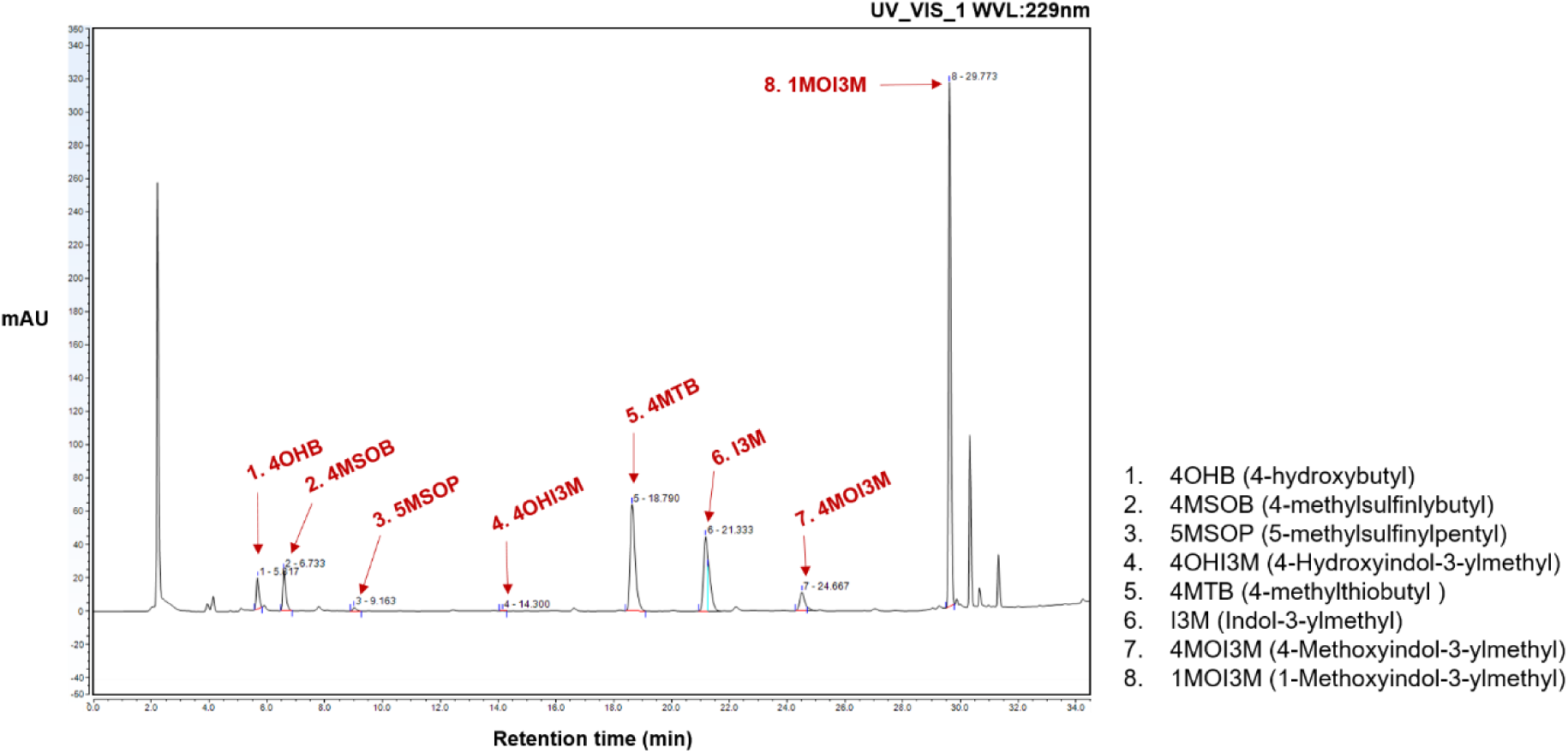
HPLC chromatogram of one-week-old *Arabidopsis* seedlings. Two-week-old seedlings grown under long day (16h light: 8h dark) at 22℃ were harvested at ZT4 and subjected for HPLC analysis (n =3). A total of eight GSL compounds (four aliphatic and four indolic GSLs) were detected using our HPLC detection system. Individual peak of GSL compound was indicated with red color arrow. Abbreviations: 4OHB, 4-hydroxybutyl; 4MSOB, 4-methylsulfinylbutyl; 5MSOP, 5-methylsulfinylpentyl; 4MTB, 4-methylthiobutyl; 4OHI3M, 4-hydroxyindol-3-ylmethyl; I3M, indol-3-ylmethyl; 4MOI3M, 4-methoxyindol-3-ylmethyl; 1MOI3M, 1-methoxyindol-3-ylmethyl. The X-axis represents retention time (minutes), and the Y-axis represents absorbance (mAU).

**Supplementary Fig. S2.**
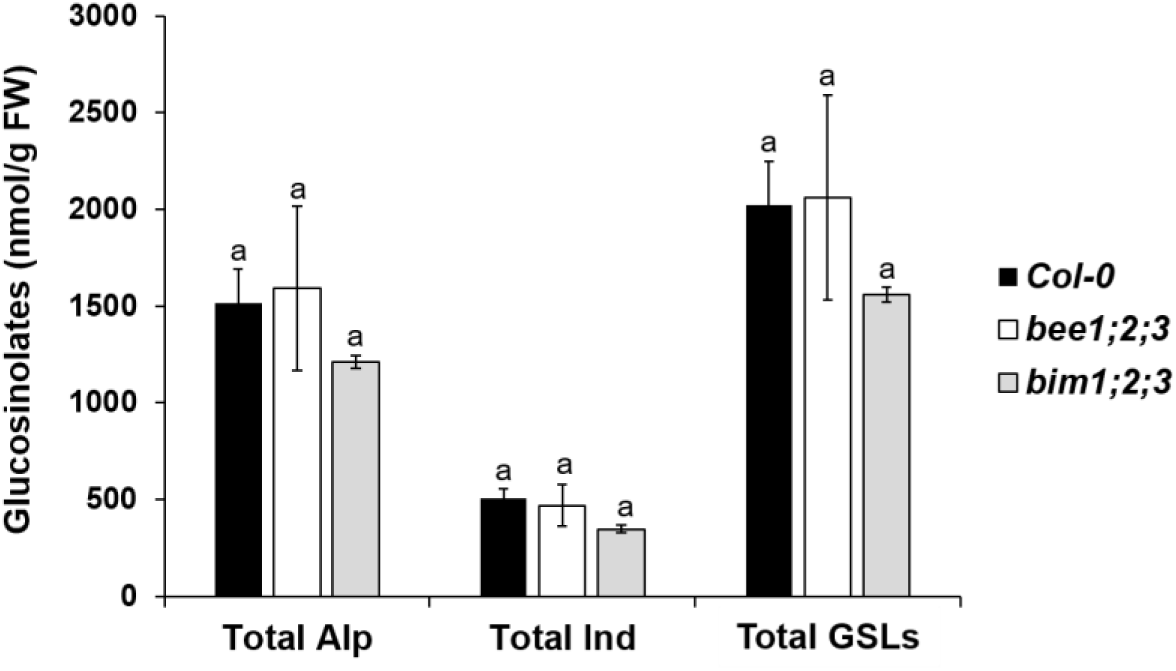
Result of HPLC quantification of aliphatic and indolic GSL compounds between Col-0, *bee1;bee2;bee3* (*bee1;2;3*), *bim1;bim2;bim3* (*bim1;2;3*) triple mutants. Amounts of total GSLs were calculated by combining amounts of aliphatic and indolic GSL compounds. Two-week-old seedlings grown under long day (16h light: 8h dark) at 22℃ were harvested at ZT4 and subjected for HPLC analysis. One-way ANOVA with Tukey’s post-hoc test was applied to calculate the statistical significance, and significant difference was indicated in the figures by different letters (p < 0.05). Data were presented as means ± standard deviation (SD) of three biological replicates. Alp: aliphatic, Ind: indolic.

**Supplementary Fig. S3.**
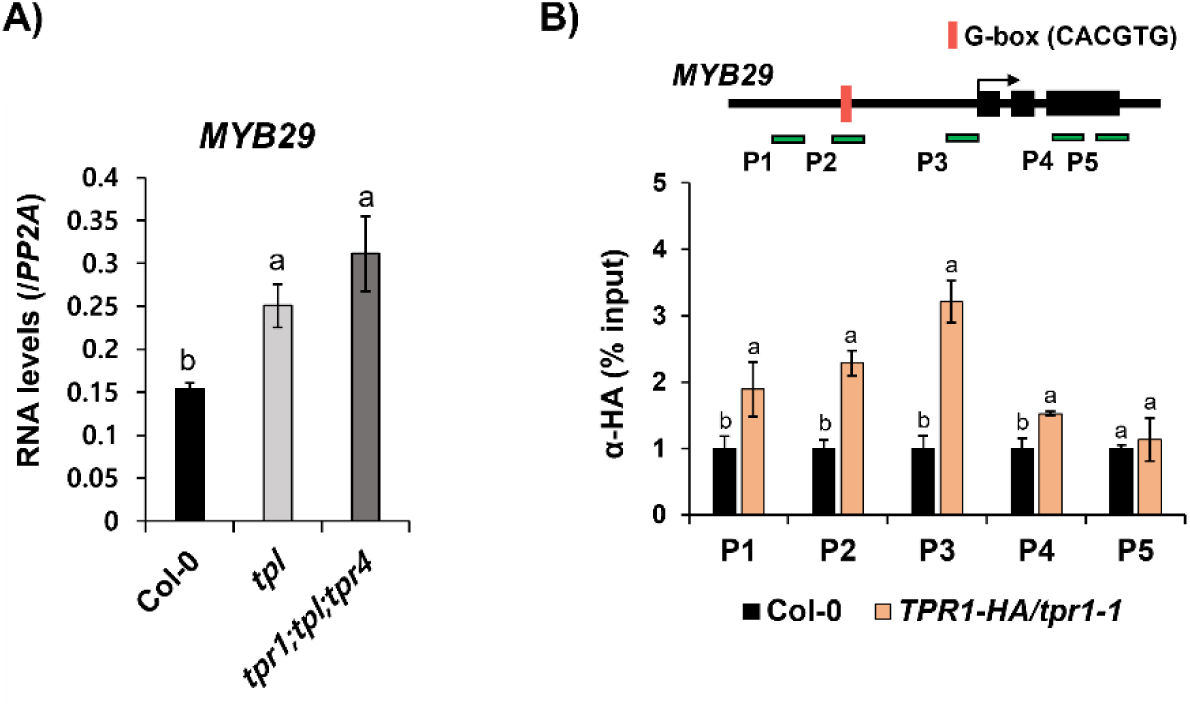
TPL and its paralogs, TPR1 and TPR4 acts redundantly on suppression of *MYB29*. **A)** Result of RT-pPCR analysis on *MYB29* between Col-0, *tpl*, and *tpl;tpr1;tpr4* mutants. **B)** Result of ChIP-PCR analysis using anti-HA antibody (ab9110, Abcam, UK) on genomic region of *MYB29* between *Col-0* and *pTPR1::TPR1-HA/tpr1-1* transgenic line. Genomic structure of *MYB29* gene and relative location of PCR amplicons used for ChIP-qPCR analysis (indicated with PX, X is arabic number) were shown above each bar graph. Eriched level of individual PCR amplicon in the Col-0 was converted to 1, and then relative enriched level of individual PCR amplicon of the *pTPR1::TPR1-HA/tpr1-1* were indicated. Two-week-old seedlings grown under long day (16h light: 8h dark) at 22℃ were harvested at ZT4 and subjected for HPLC analysis. **A)∼B)** One-way ANOVA with Tukey’s post-hoc test was applied to calculate the statistical significance, and significant difference was indicated in the figures by different letters (p < 0.05). Data were presented as means ± standard deviation (SD) of three biological replicates.

**Supplementary Fig. S4.**
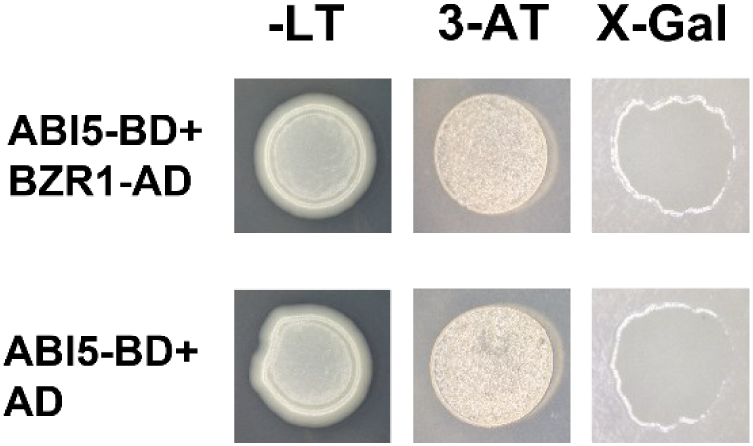
Result of yeast 2 hybrid (Y2H) assay between ABI5 and BZR1 or BES1. ABI5 (bait) and BZR1 (prey) exhibited no interaction in yeast two hybrid (Y2H) assay between bait and prey. –LT: a double-dropout SD (synthetic defined) media lacking leucine and tryptophan. 3AT: 3-Amino-1, 2, 4-triazole (3-AT) (20 mM). X-Gal: X-alpha-gal assay.

**Supplementary Fig. S5.**
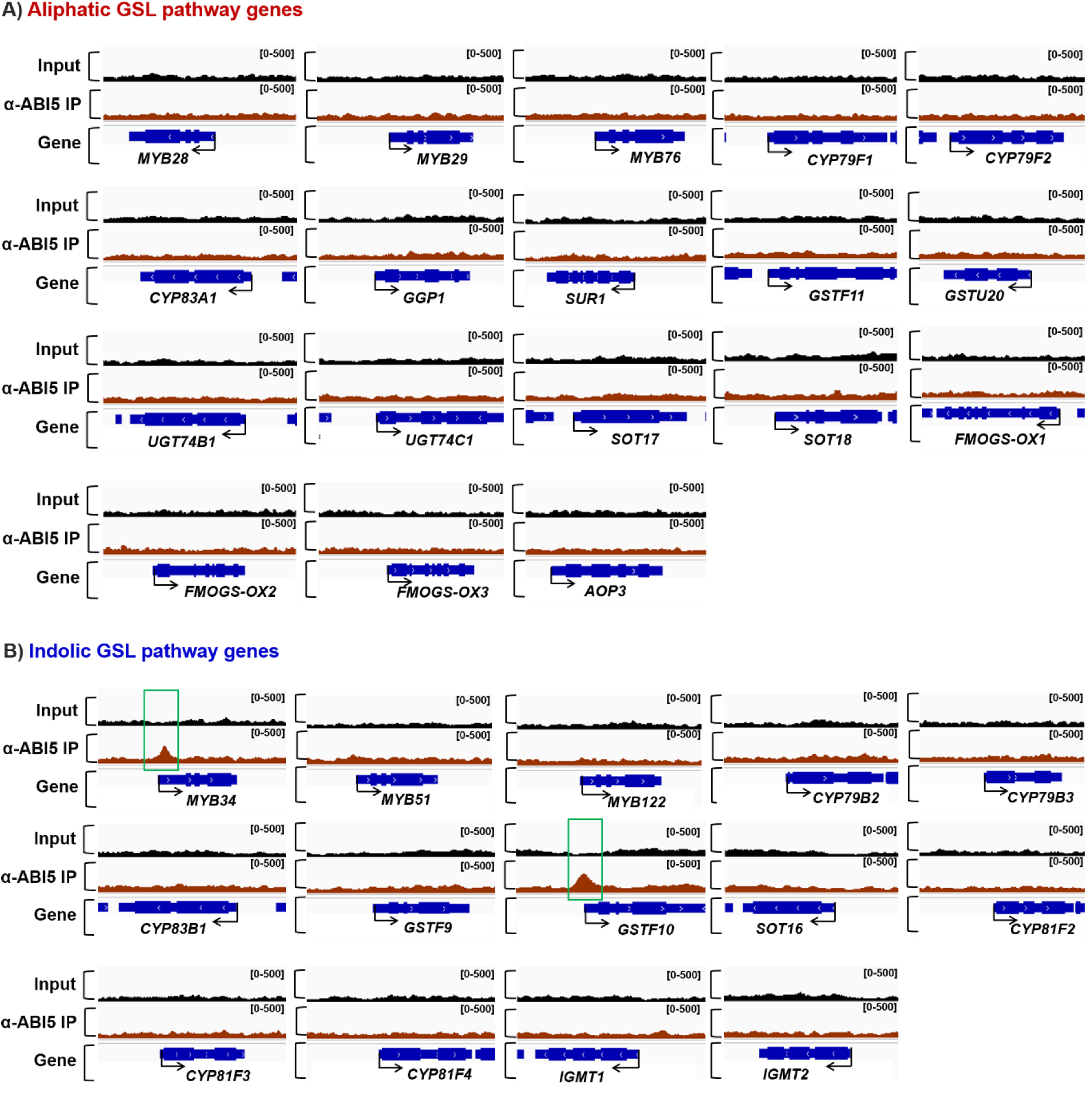
Integrative genome viewer (IGV) illustration of ChIP-seq dataset using α-ABI5 on GSL pathway genes. **A)** IGV illustration of ChIP-seq dataset using α-ABI5 on aliphatic GSL pathway genes. No significant binding of ABI5 was detected in any of aliphatic GSL pathway genes in this dataset. **B)** IGV illustration of ChIP-seq dataset using α-ABI5 on indolic GSL pathway genes. Among indolic GSL pathway genes, only *MYB34* and *GSTF10* were found to have a significant peak of ABI5 at around transcription start region of these genes. Significant binding of ABI5 on *MYB34* and *GSTF10* was indicated with green box. **A)∼B)** Number in the bracket at the upper right corner shows the range of normalized read counts mapped to the annotated individual genes. The black and red color tracks represent input DNA and α-ABI5-immunoprecipitated DNA, respectively. Gene structures (blue color) and gene names were shown at the bottom of the each IGV browser view. Dataset showing ABI5 enrichments on aliphatic GSL pathway genes (A) and indolic GSL pathway genes (B) were generated with α-ABI5 antibody (ab98831, Abcam, UK). Detailed information on this public available ChIP-seq dataset is provided in Supplementary Table S3.

**Supplementary Fig. S6.**
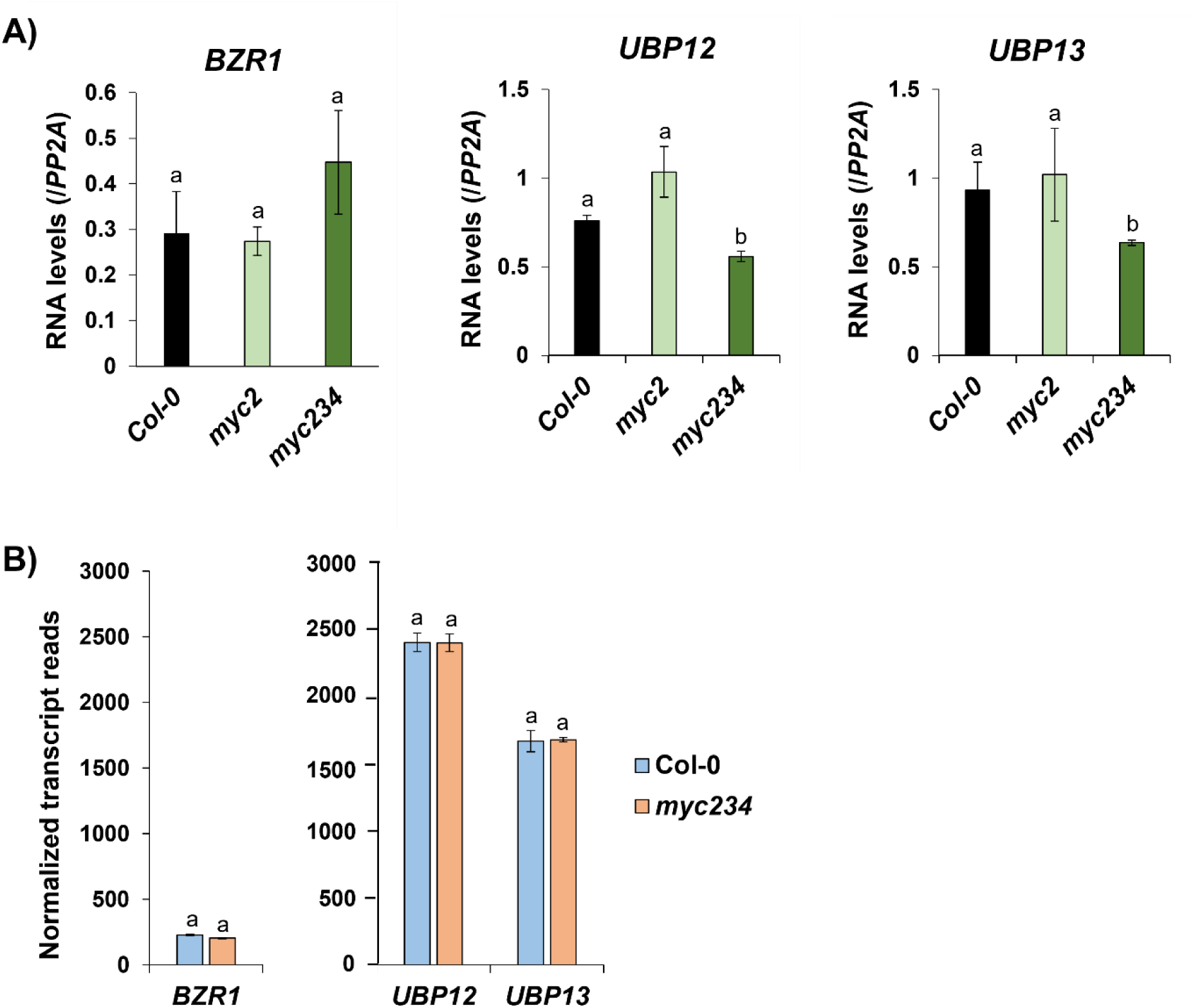
Transcript levels of *BZR1, BES1, UBP12,* and *UBP13* genes between Col-0 and *myc2;3;4* mutant. **A)** Transcript levels of *BZR1, UBP12,* and *UBP13* genes between Col-0, *myc2* single mutant, and *myc2;3;4* triple mutant. Two-week-old seedlings grown under long day (16h light: 8h dark) at 22℃ were harvested at ZT4 and subjected for RT-qPCR analysis (n =3). One-way ANOVA with Tukey’s post-hoc test was applied to calculate the statistical significance, and significant difference was indicated in the figures by different letters (p < 0.05). **B)** Normalized transcript levels of *BZR1, UBP12,* and *UBP13* genes between Col-0 and *myc2;3;4* mutants. A public RNA-seq dataset (Van Moerkercke et al., 2019) was analyzed to obtain normalized transcript levels of those four BR signaling genes. Information on the RNA-seq dataset used for this study was shown in the Supplementary Table S3.

**Supplemental Table S1.**
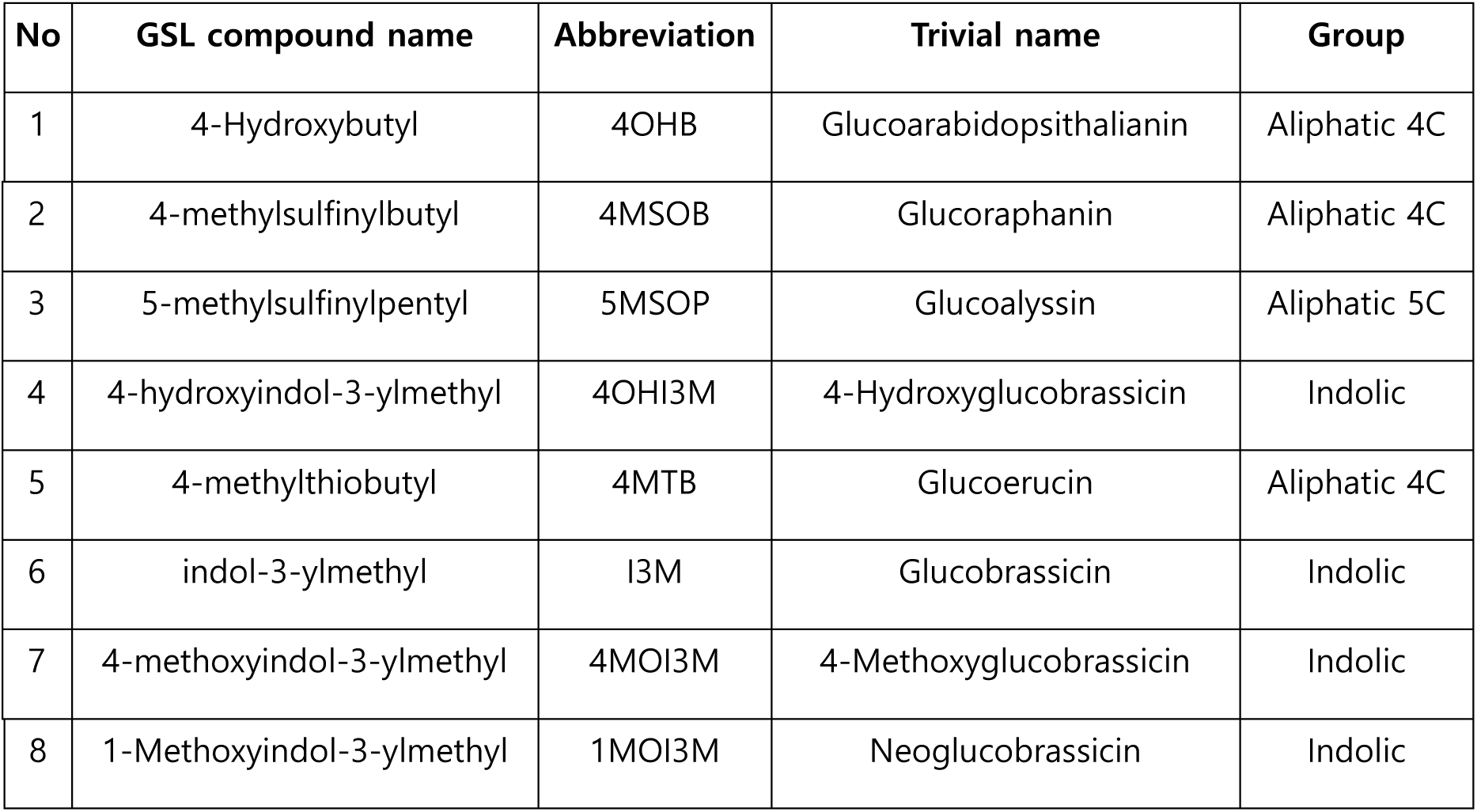
Major GSLs identified by LC-ESI/MS from two-week-old *Arabidopsis* seedlings.

**Supplemental Table S2.**
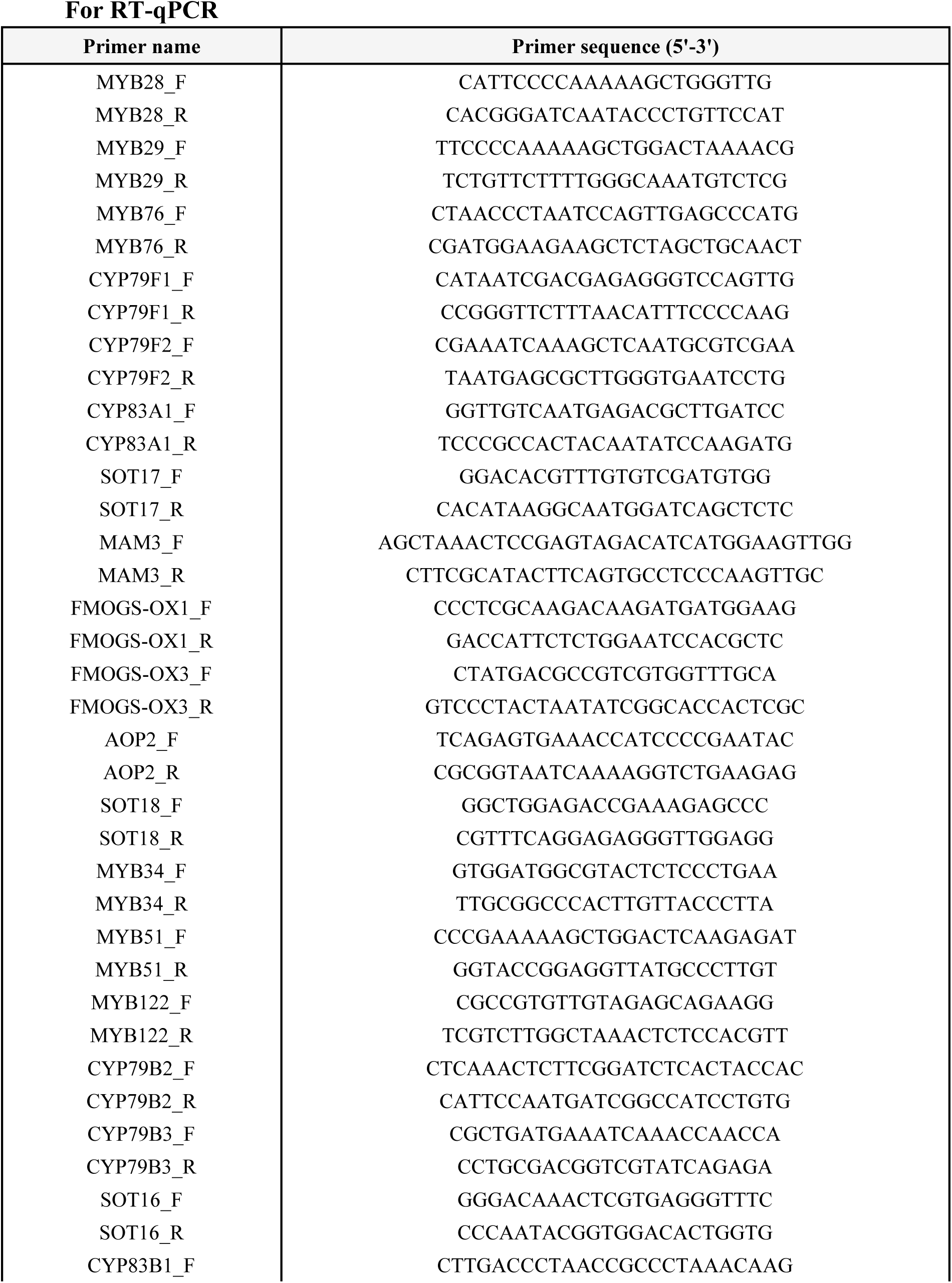
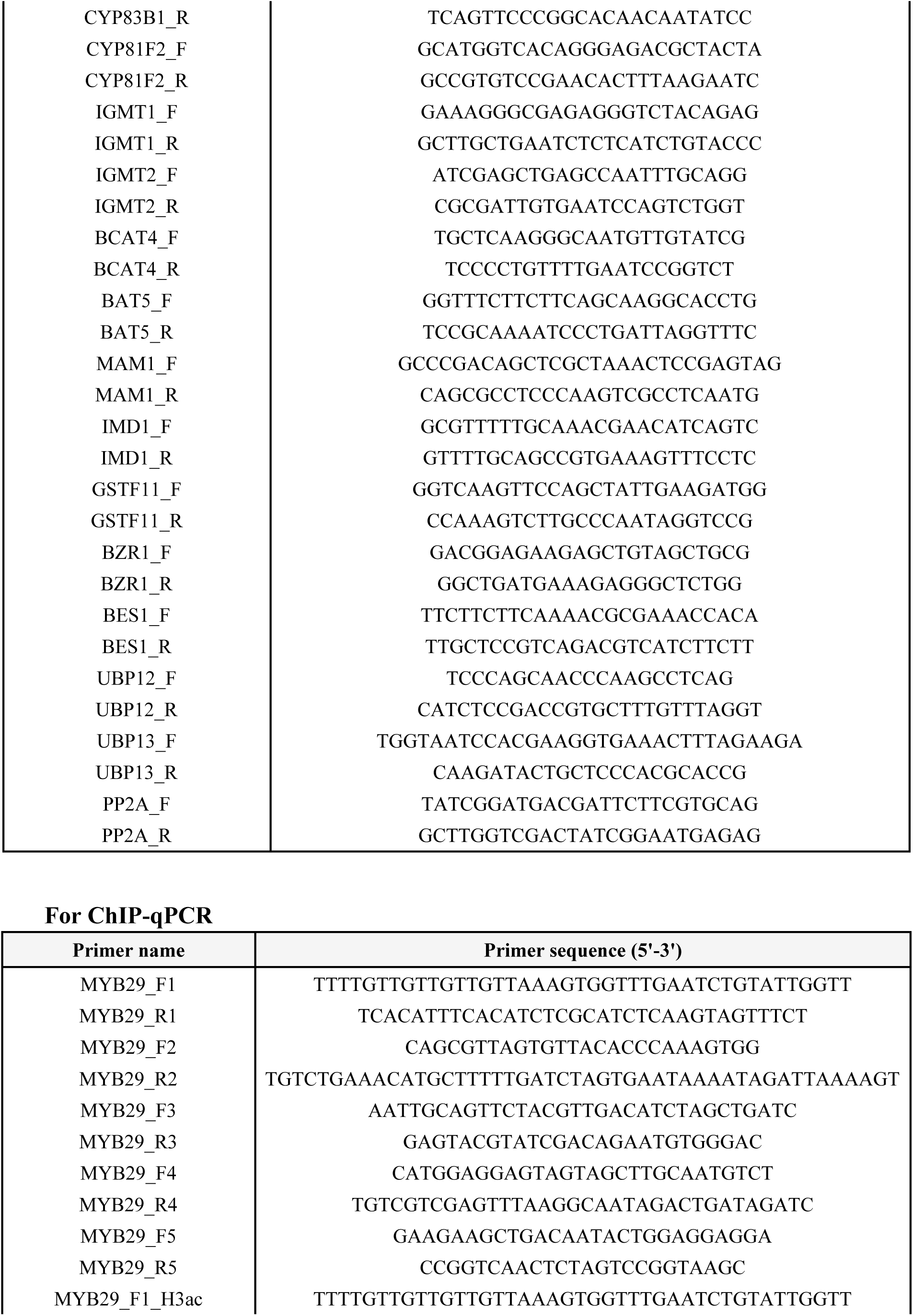
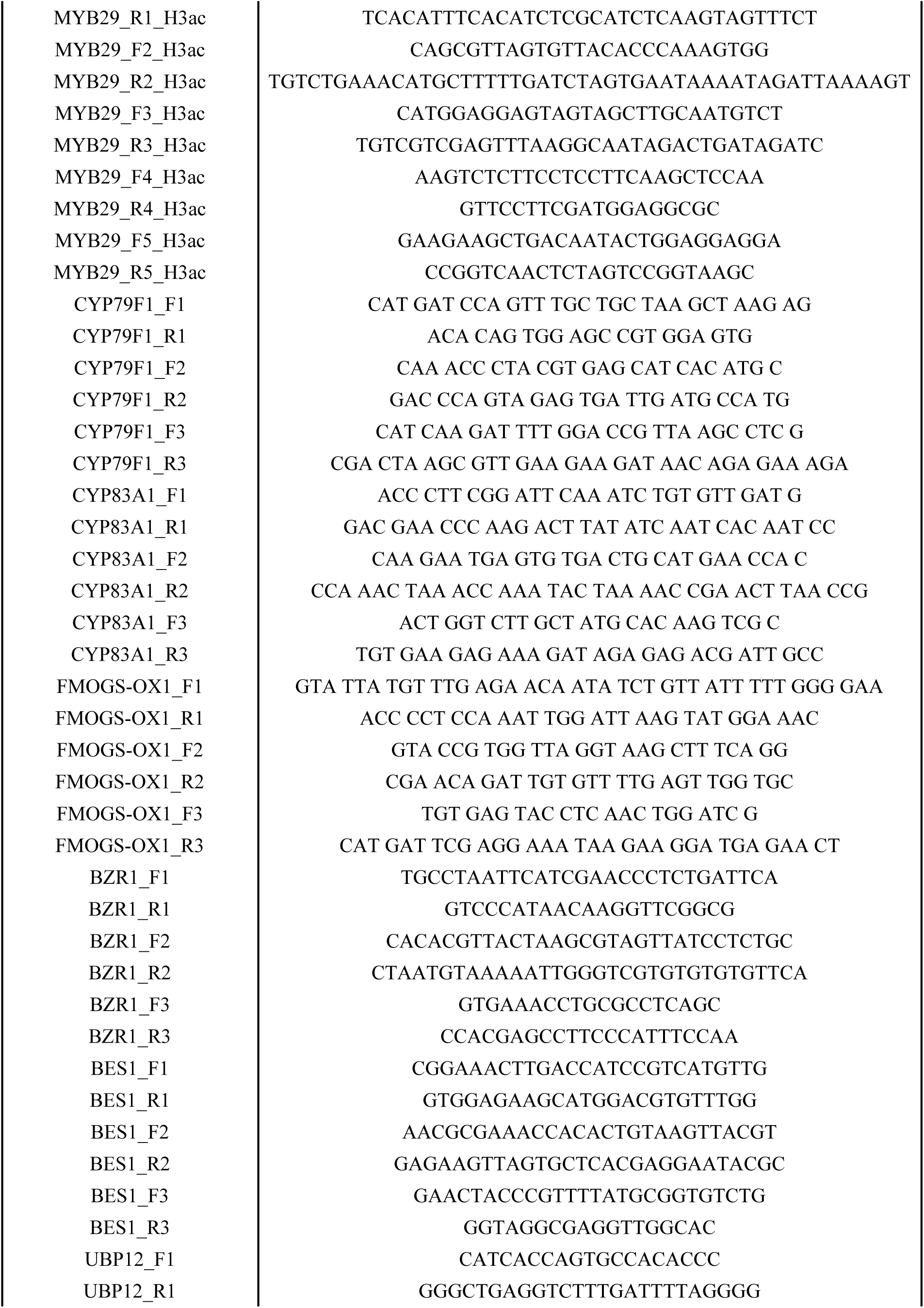
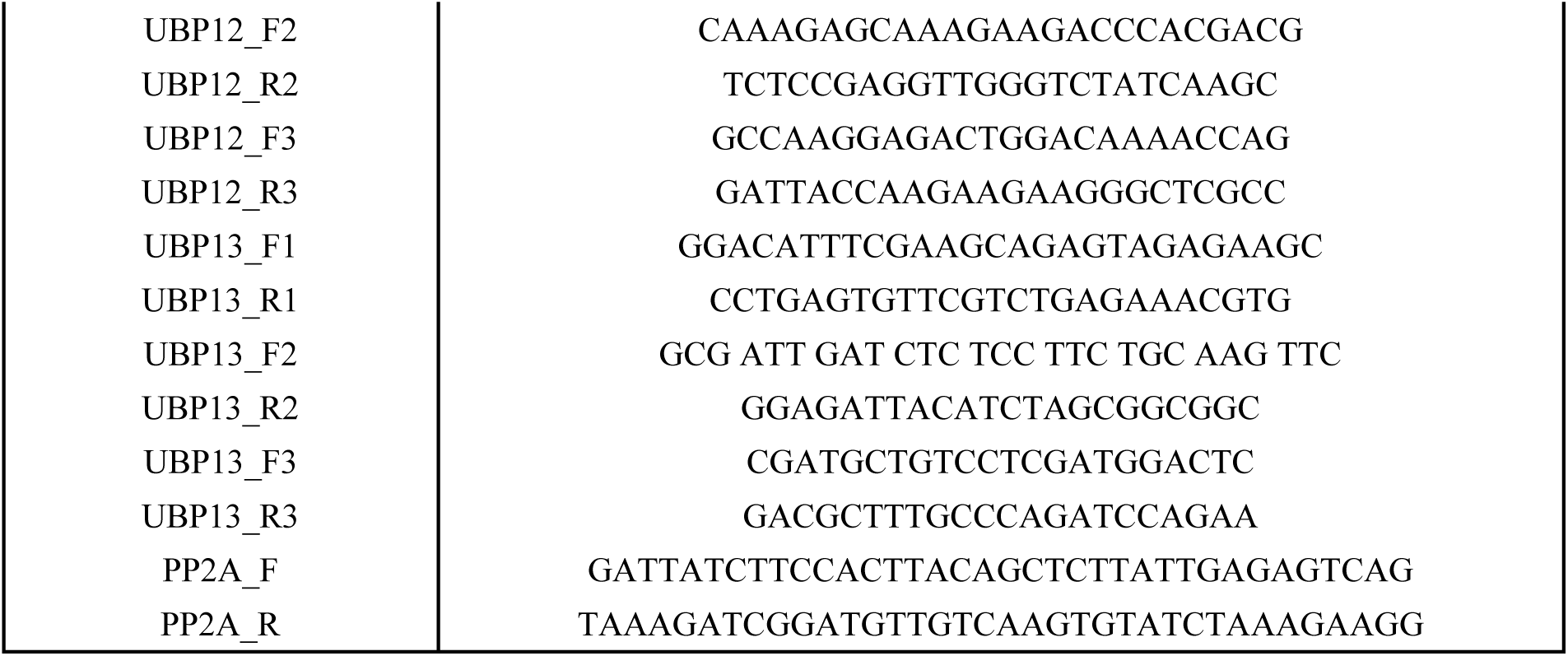
Information on the primers used in the RT-qPCR and ChIP-qPCR assay in this study.

**Supplementary Table S3.** Information of public RNA-seq and ChIP-seq dataset analyzed in this study.

| Type | Samples | Accession number | Database |
| --- | --- | --- | --- |
| RNA-seq | Col-0 rep 1 | E-MTAB-8019 | ArrayExpress database |
|  | Col-0 rep 2 |  |  |
|  | Col-0 rep 3 |  |  |
|  | *myc2;3;4* rep 1 |  |  |
|  | *myc2;3;4* rep 2 |  |  |
|  | *myc2;3;4* rep 3 |  |  |
| ChIP-seq | Col-0 rep 1 | SRR13619483 | EMBL-EBI database |
|  | Col-0 rep 2 | SRR13619484 |  |
| | α-ABI5 IP rep 1 | SRR13619490 | |
| | α-ABI5 IP rep 2 | SRR13619491 | |

